# Twitching motility suppressors reveal a role for FimX in type IV pilus extension dynamics

**DOI:** 10.1101/2025.07.10.664058

**Authors:** Nathan Roberge, Nathan Yuen, Hanjeong Harvey, Taylor J. Ellison, Courtney K. Ellison, Lori L. Burrows

## Abstract

In *Pseudomonas aeruginosa,* retractable protein filaments called type IV pili (T4P) facilitate surface adherence, sensing, and directional movement known as twitching motility. T4P are necessary for the bacteria to engage in surface-associated behaviors, including establishing acute infections. Pilus extension is driven by the hexameric ATPase, PilB, at the base of the T4P nanomachine in coordination with various protein regulatory effectors. The cyclic-di-GMP binding protein, FimX, works with PilB to mediate normal extension processes, though how this effector controls pilus assembly remains unclear. To explore the role of FimX in T4P function, we leveraged the significant Δ*fimX* twitching motility deficit to screen for mutants capable of overcoming this phenotype. We identified suppressor mutations that increase twitching in Δ*fimX* background, mapping primarily to cyclic-AMP homeostatic machinery or to PilB, the FimX target. Distinct suppressor mutations in PilB increased ATP hydrolysis *in vitro* and this activity was subject to modulation by FimX. Using microscopy to monitor the extension dynamics of fluorescently labelled T4P, we showed that Δ*fimX* mutants produce slow-to-extend, short pili, a phenotype that is rescued by mutations enhancing PilB ATP hydrolysis and/or re-introduction of FimX. Together, these data implicate FimX as a regulator of PilB enzymatic function, potentially enabling *P. aeruginosa* to fine-tune pilus extension dynamics in response to environmental cues.

**Summary:** Type IV pili enable *Pseudomonas aeruginosa* to attach to surfaces, move (twitch), and form biofilms. Pilus extension is powered by the motor protein PilB, which is regulated by other factors, including FimX, a protein that binds cyclic-di-GMP. Although FimX is important for twitching, how it influences PilB was unclear. We deleted *fimX*, which severely reduces motility, and searched for mutants that regained movement. We identified two types: some had mutations in PilB that increased its ATPase activity, allowing it to function without FimX, while others affected the cyclic-AMP signaling pathway and increased overall production of pilus components, showing that motility can also be improved through changes in quantity versus quality. Our results suggest that FimX normally fine-tunes PilB enzymatic activity, enabling dynamic control of pilus extension in response to surface signals. This work helps explain how *P. aeruginosa* adapts to different environments, a process crucial for infection and biofilm development.

## Introduction

*Pseudomonas aeruginosa* deploys type IV pili (T4P) to adhere to and spread across surfaces (1, 2). Directed bacterial movement, or twitching motility, is powered by repeated cycles of protein filament extension, surface attachment, and retraction to pull cells along a surface (3–5). Pilus function is coordinated through the action of four distinct protein subcomplexes which together comprise the T4P nanomachine (Fig 1A) (6). T4P also allow cells to sense surface contact and assist in the transition from planktonic to sessile lifestyles (5, 7–9), facilitating microcolony formation, and eventually, biofilm production (10, 11). Engagement of the extracellular filament with a surface induces an as-yet undefined signal that is propagated to the inner membrane embedded Pil-Chp regulatory network, which in turn stimulates CyaB-dependent synthesis of the secondary messenger molecule, cyclic adenosine monophosphate (cAMP) (Fig 1A) (5, 7, 12). High intracellular cAMP activates the transcription factor Vfr to modulate the expression of over 300 genes, including those encoding T4P machinery components (13), with cAMP homeostasis maintained by the action of the CpdA phosphodiesterase (14–16). The cascade of secondary messenger signaling culminates in elevated cyclic diguanylate monophosphate (cdGMP) (16), which in many bacteria is associated with sessile phenotypes such as biofilm formation (17). This increase in cdGMP potentially occurs through stimulation of T4P-associated diguanylate cyclases (DGCs) like SadC (18, 19), and/or through surface sensing by pilus tip-associated proteins (6, 16, 20, 21).

**Figure 1:**
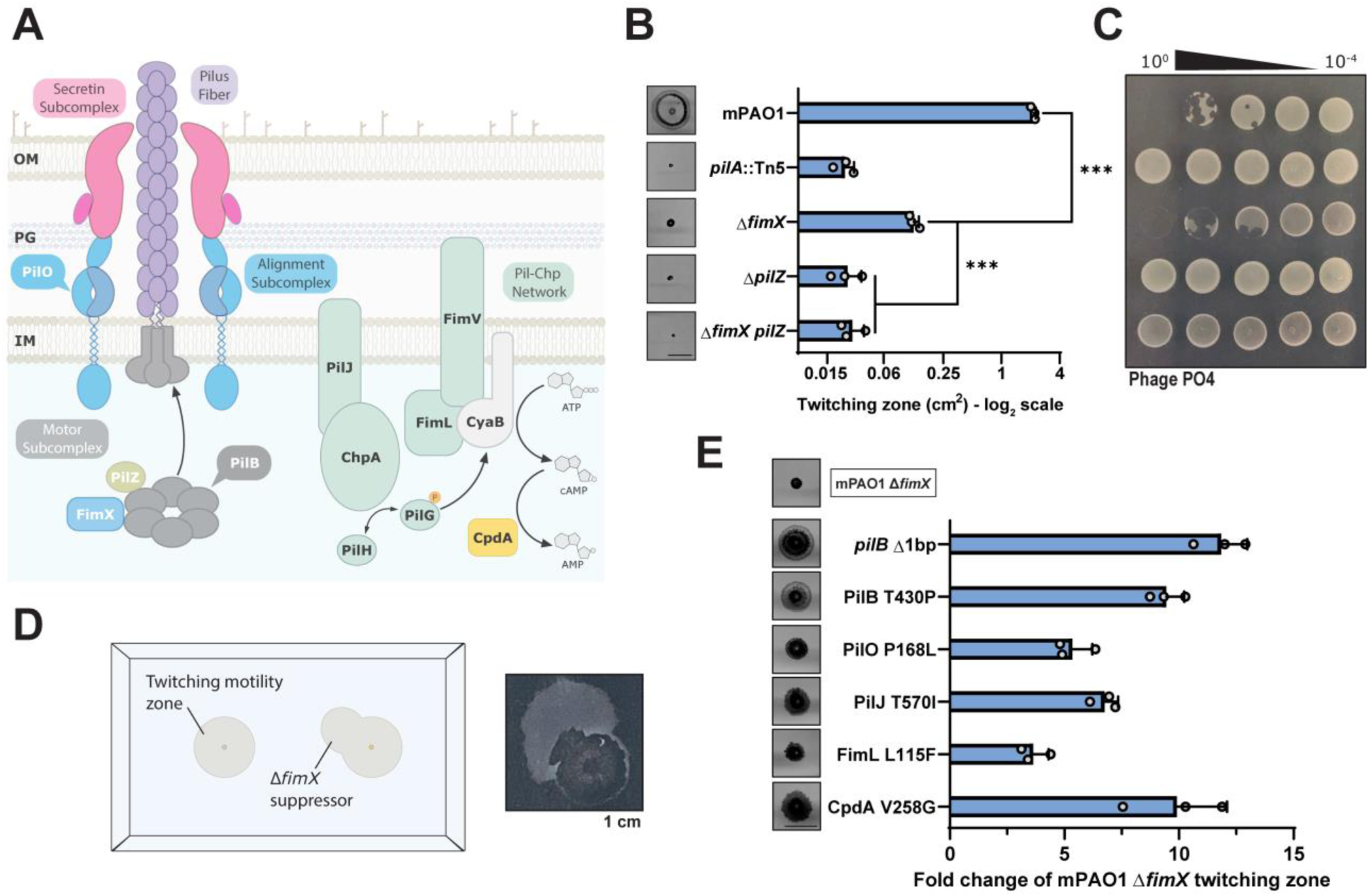
Suppressor mutations overcome Δ*fimX* twitching motility defects. **(A)** The type IV pilus nanomachine comprising four distinct protein subcomplexes (motor, alignment, secretin subcomplexes, and pilus fiber) and associated inner-membrane Pil-Chp signaling network responsible for cAMP generation. PilB in association with regulatory effectors PilZ and FimX drives pilus extension. **(B)** Quantification of sub-agar stab twitching motility zones for extension regulatory effector mutants. Representative crystal violet-stained twitching zones are shown to the left. X-axis depicts a log_2_ scale. Bars represent the means of triplicate samples from three independent experiments ± SD. **(C)** Representative plaque assay demonstrating 10-fold serial dilutions of bacteriophage PO4 on extension regulatory effector mutants. **(D)** Cartoon representation of the Δ*fimX* suppressor mutant screen setup. A representative motile flare is shown to the right. Image border = 1 cm. **(E)** Quantification of sub-agar stab twitching motility zones for selected Δ*fimX* suppressor mutants. Representative twitching zones are shown to the left. Data are plotted as a fold change of the Δ*fimX* parent strain. Bars represent the means of triplicate samples from three independent experiments ± SD. All scale bars = 1 cm. **\*\*\***: *p* ≤ 0.001 (Two-tailed parametric *t*-test).

Pilus extension is an essential first step in filament biogenesis. Extended pili can interact with external substrates, ultimately triggering downstream signaling and facilitating motility (2, 8, 22, 23). Extension is driven by the hexameric ATPase PilB (1, 24–26) in conjunction with regulatory effectors PilZ (27, 28) and the cdGMP-binding protein FimX (27, 29–31) (Fig 1A), both of which are necessary for normal pilus assembly. The role of FimX in pilus extension is not well understood, with studies of *P. aeruginosa* FimX and its homologues in other species implicating a variety of disparate functions (27, 30, 32, 33). In *P. aeruginosa*, FimX interaction with PilB is contingent upon cdGMP binding by FimX’s EVL motif in the C-terminal EAL domain. *P. aeruginosa* FimX mutants that are unable to bind the secondary messenger are defective in surface piliation (30, 31), twitching motility (29–31), and ATPase localization (27, 30). Some of these phenotypes can be overcome through suppressor mutations that increase cellular cdGMP levels. Those studies suggest that FimX is responsible for maintaining PilB functional efficiency over broad concentrations of cdGMP (31). On the other hand, more recent work showed selective trafficking of the PilB-FimX subcomplex to the leading poles of twitching cells. This localization promotes biased pilus extension at a single pole, resulting in increased Pil-Chp network signaling at the same pole. The positive feedback loop was proposed to ensure directional movement of the rod-shaped cells in response to PilB localization. Further, PilB polar localization was dependent upon FimX and various cytoplasmic components of the Pil-Chp network, suggesting that FimX may also be a cdGMP-binding polar localization factor that coordinates optimal PilB subcellular distribution (8, 34).

How then, do these proposed functions intersect and what is the role of FimX in *P. aeruginosa* pilus function and/or signaling? To shed light on how FimX controls pilus function, we leveraged the Δ*fimX* twitching motility deficit to select for suppressor mutants capable of overcoming loss of its input. Our screen revealed two classes of mutants with increased motility in the absence of FimX. These mutations were primarily in the Pil-Chp network responsible for modulating cAMP levels or PilB, the interaction partner of FimX. We focused on the PilB mutations, which notably were not clustered but rather mapped to various regions of the protein. We characterized two distinct examples and demonstrated that each significantly increased PilB ATP hydrolysis *in vitro*. Phenotypically, this elevated activity translated to increased rates of extension, resulting in longer pili. Both ATPase activity and pilus extension rates were subject to FimX-dependent modulation. Therefore, FimX modulates PilB enzymatic efficiency to produce longer pili suited for twitching motility.

## Results

### Suppressor mutations restore motility in Δ*fimX* mutants

FimX is a regulator of pilus extension in *P. aeruginosa*, though its exact mechanism is not well understood (29–31). Consistent with previous work, we confirmed that a PAO1 deletion mutant of *fimX* had significantly reduced twitching motility, 5-10% of the wild type (Fig 1B). However, this strain still twitched when compared to a non-motile major pilin (*pilA*::Tn5) mutant and remained susceptible to the pilus-targeting bacteriophage, PO4 (Fig 1C). FimX mutants did not produce detectable surface pili in sheared protein fractions (Fig S1), further supporting a defect in pilus extension. This phenotype is distinct from that of mutants lacking another PilB regulator, the putative chaperone, PilZ (27, 28), which do not twitch and are resistant to pilus-specific phages (Fig 1B and C).

We reasoned that we could leverage the Δ*fimX* pilus extension deficit to screen for suppressor mutants capable of restoring twitching motility, to inform on the function of FimX. After allowing *fimX* mutants to twitch at the sub-agar interface for an extended period (48-72 h), we isolated highly motile flares that emerged from the small primary twitching zone (Fig 1D). These cells were collected, isolated for single colonies, and after confirming their increased motility compared to the Δ*fimX* parent, were sequenced to identify candidate suppressor mutations. The mutations mapped to relevant proteins including the ATPase PilB, alignment subcomplex component PilO (35, 36), members of the Pil-Chp network (PilJ and FimL) (7), and CpdA (additional mutants are listed in Table S1). Each mutation increased motility to varying degrees (Fig 1E), though not to wild-type levels, but did not impact growth (Fig S2) or phage susceptibility (Fig S3). Complementing the suppressor mutants with *fimX* further increased twitching motility to levels similar to or greater than WT, except for PilO P168L, suggesting it has a *fimX* independent motility phenotype (Fig S4A). In contrast, complementation of the suppressors with a defective version of FimX – where its cdGMP-binding EVL motif was mutated to AAA – did not further increase twitching motility, consistent with previous reports that this non-functional variant is incapable of PilB binding (30). Diminished twitching motility in the WT strain when FimX AAA was expressed in trans is likely due to formation of FimX-FimX AAA heterodimers, supported by their pairwise interactions in a bacterial 2-hybrid assay (Fig S4B).

### Δ*fimX* suppressor mutations do not directly influence cdGMP metabolism

Previous efforts to isolate Δ*fimX* suppressors revealed several mutations which collectively upregulated intracellular cdGMP levels (31). To test whether our mutants shared this phenotype, we assessed intracellular levels of cdGMP using a *pcdrA*:Lux cassette-based reporter system in liquid cultures (20). Under these conditions, none of our mutants demonstrated higher cdGMP levels over time when compared to control cells expressing the well-characterized diguanylate cyclase (DGC), SadC (Fig 2A). To directly test the impact of cdGMP on Δ*fimX* suppression, we overexpressed DGCs from *P. aeruginosa* (*sadC*) or *Escherichia coli* (*ydeH*) in WT or Δ*fimX* backgrounds and tested twitching motility. In all cases, cells expressing the DGCs showed significantly reduced twitching motility compared to the vector control (Fig 2B). Together, these data suggest that suppression of the Δ*fimX* motility defect in PAO1 is not related to changes in cdGMP levels.

**Figure 2:**
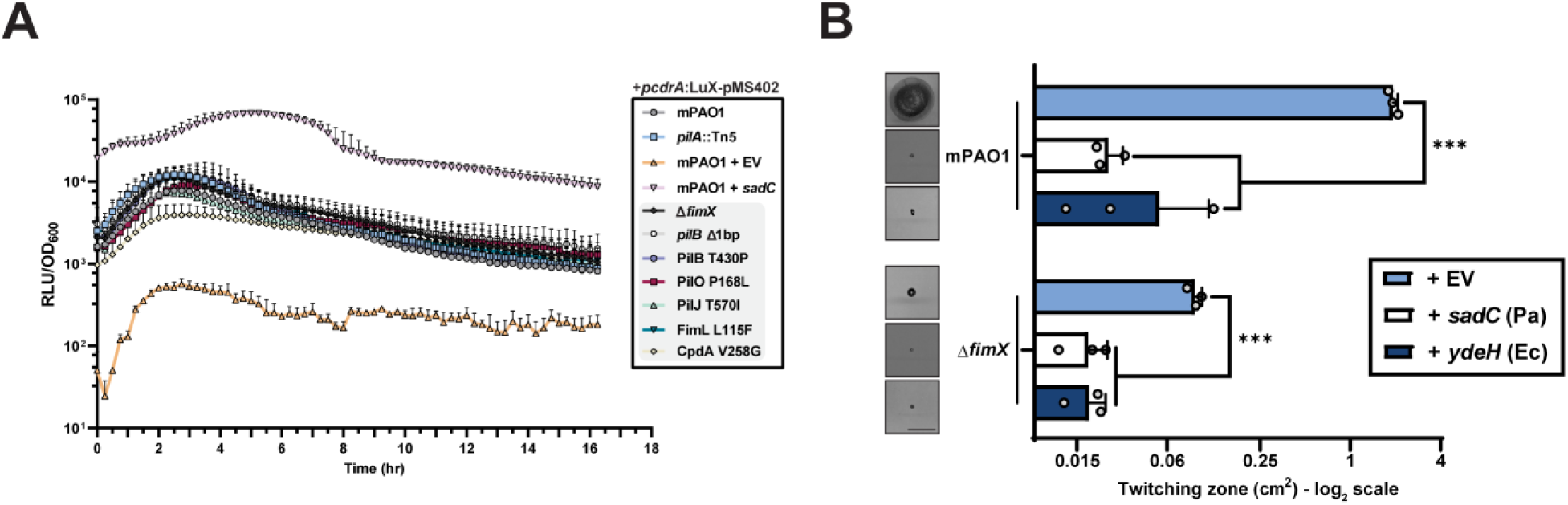
FimX suppressor mutations do not impact intracellular cdGMP levels. **(A)** Time-course curves of relative luminesence signal of a *pcdrA*:LuX cassette cdGMP reporter normalized to OD_600_. Each timepoint is the mean of triplicate samples from three independent experiments ± SD. **EV**: empty pMS402 vector, ***sadC***: *sadC* in pBADGr vector. **(B)** Quantification of sub-agar stab twitching motility zones for strains overexpressing diguanylate cyclases (DGCs). Representative crystal violet-stained twitching zones are shown to the left. DGC expression was induced with 0.1% arabinose. Bars represent the means of triplicate samples from three independent experiments ± SD. X-axis depicts a log_2_ scale. Scale bar = 1 cm. **EV**: empty pBADGr vector, ***sadC***: *sadC* in pBADGr vector, ***ydeH***: *ydeH* in pBADGr vector. **\*\*\***: 0.001 ≥ *p* (Two-tailed parametric *t*-test).

### Elevated cAMP levels increase motility in the Δ*fimX* background

Since several of our motile suppressor mutations mapped to the Pil-Chp network or CpdA, we hypothesized that changes to cAMP homeostasis (5, 7) increased twitching in the absence of *fimX*. We therefore assessed intracellular levels of cAMP in both liquid cultures and surface-adapted cellular populations using a *PaQa*:YFP reporter plasmid system (37). WT cells had significantly higher levels of cAMP when the cells were grown on a surface versus in liquid, but elevated cAMP-dependent signal was lost in a *pilA* mutant (Fig 3A), consistent with T4P being a major input to cAMP generation in *P. aeruginosa* (38). Deletion of CyaB or expression of a constitutively active CyaB R456L *in trans* further reduced or increased cAMP levels, respectively (15, 39, 40). In these assay conditions, Δ*fimX* had lower than WT levels of cAMP and was unable to respond to surface contact. The Δ*fimX* suppressor mutants PilJ T570I, FimL L115F, and CpdA V258G all had significantly elevated cAMP levels. In contrast, neither the PilB nor PilO P168L suppressor mutants had significant increases in cAMP levels compared to the Δ*fimX* parent. These data suggested that there were at least two routes to increase twitching in the *fimX* background, one of which was independent of cAMP levels.

**Figure 3:**
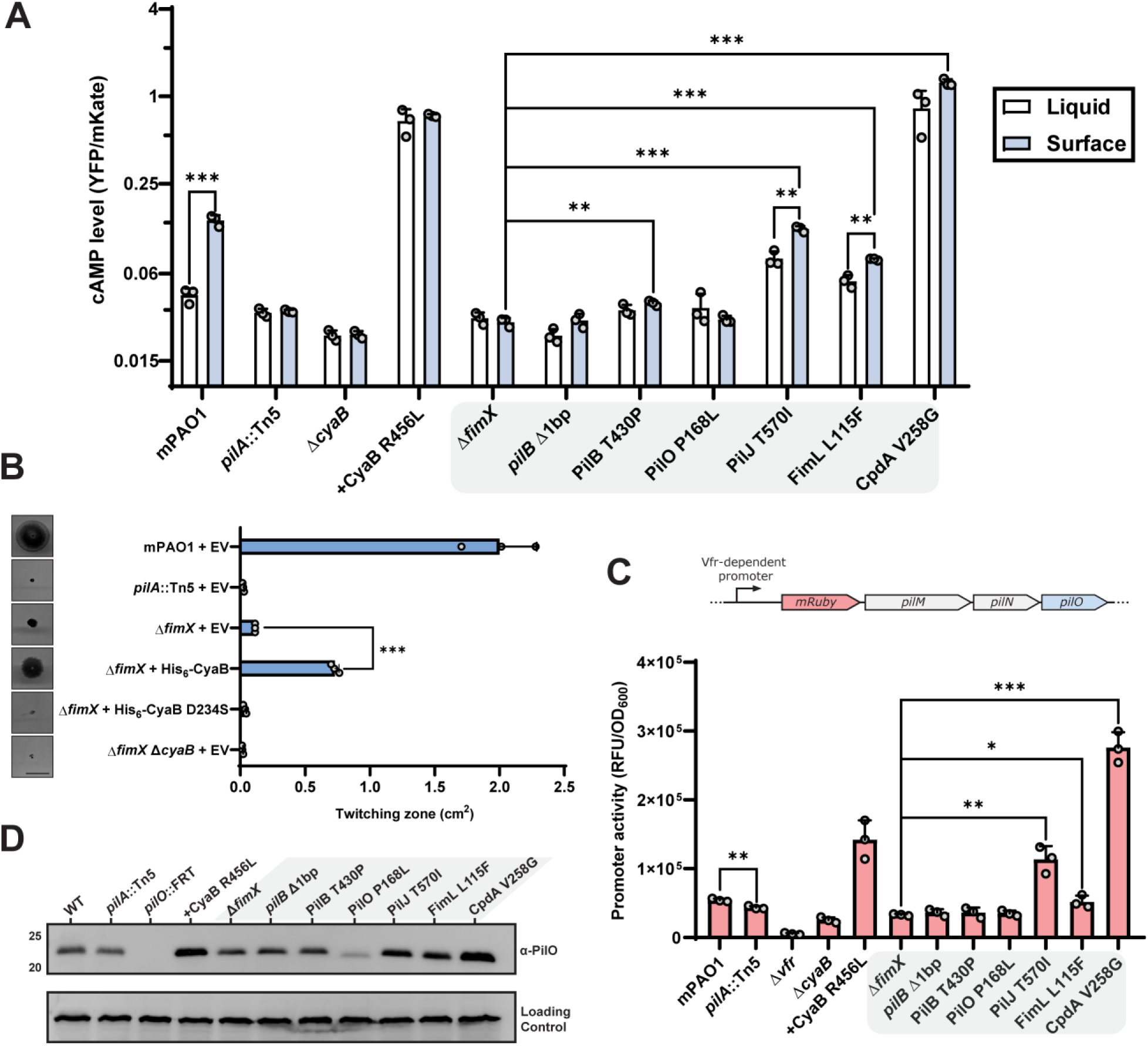
A subset of Δ*fimX* twitching suppressors elevate cAMP levels and upregulate associated phenotypes. **(A)** Plasmid-based cAMP reporter assay quantifying cAMP-dependent YFP expression (*PaQa*) normalized to constitutive mKate expression (*rpoD*). Bars represent the means of duplicate samples from three independent experiments ± SD. **(B)** Quantification of sub-agar stab twitching motility zones for strains overexpressing the adenylate cyclase CyaB or a catalytically inactive mutant. Representative crystal violet-stained twitching zones are shown to the left. Expression was induced with 0.1% arabinose. Bars represent the means of triplicate samples from three independent experiments ± SD. Scale bar = 1 cm. **EV**: empty pHERD30T vector, **His_6_-CyaB (D234S)**: N-terminally hexa-histidine tagged CyaB or CyaB D234S in pHERD30T vector. **(C)** Vfr-dependent expression of mRuby3 chromosomally inserted upstream of the native alignment complex *pilMNOPQ* operon. Bars represent the means of duplicate samples from three independent experiments ± SD. Cartoon schematic of the mRuby3 expression system is shown above the graph. **(D)** Representative PilO Western blot. Sample loading was normalized to a non-specific band shown below. Molecular weight is indicated to the left. Blot is representative of three independent experiments. **CyaB R456L**: CyaB R456L in pBADGr vector. **\***: 0.05 ≥ *p* ≥ 0.01; **\*\***: 0.01 ≥ *p* ≥ 0.001; **\*\*\***: 0.001 ≥ *p* (Two-tailed parametric *t*-test).

We next asked whether high cAMP levels were responsible for increased Δ*fimX* twitching. Expressing CyaB from a multicopy plasmid in the Δ*fimX* background was sufficient to significantly increase twitching motility compared to the empty vector control (Fig 3B), while expression of an inactive CyaB variant (Fig S5A) (41) was not. This increased twitching phenotype was also observed in a double mutant lacking the Pil-Chp response regulators PilG and PilH (Fig S5B), bypassing Pil-Chp-dependent CyaB activation (34). Together, these data indicate that elevated cAMP levels due to Pil-Chp or CpdA mutations increase Δ*fimX* motility. Strains with high intracellular cAMP in turn had increased Vfr-dependent expression (13) of an mRuby3 fluorescent protein reporter integrated upstream of the T4P *pilMNOPQ* operon (Fig 3C). Integration of the reporter did not impact phage susceptibility of these strains (Fig S6) indicating that pilus function is maintained. Consistent with this increased expression, levels of PilO were also elevated (Fig 3D) suggesting that more T4P machines are expressed in these backgrounds.

Notably, the PilB and PilO suppressors had no increase in Vfr-dependent transcription or abundance of T4P components (Fig 2C and D) compared to Δ*fimX*, consistent with low cAMP levels (Fig 3A). Another cAMP-dependent trait, type II secretion system-dependent secretion of proteases generating zones of clearing on skim-milk plates (42) was also unchanged (Fig S7A and B), collectively suggesting that the cAMP levels in those mutants were not significantly impacted (13, 15), and that Δ*fimX* suppression occurs through non-cAMP related mechanisms in these backgrounds.

### Determinants for *pilB* Δ1bp function

The suppressors in the non-cAMP related class predominantly mapped to the pilus extension ATPase, PilB. To probe this second set, we focused first on a mutant in which a single adenine was deleted in the last codon of the PilB open reading frame (*pilB* Δ1bp). This deletion produced a frameshift that extended the coding sequence by an additional 21 bases, or seven amino acids, to the next stop codon (Fig 4A). When modelled in Alphfold3, the extended segments were predicted to be solvent-exposed, protruding from the base of the hexamer and away from its putative interaction interfaces with additional T4P components (Fig S8). To find the minimum length of this C-terminal extension that was sufficient to increase motility in Δ*fimX*, we generated single codon truncations from the 3ʹ end of the *pilB* Δ1bp sequence (Fig 4A). Each variant complemented motility in both a *pilB*::Tn5 background (Fig S9A), and in the WT (Fig S9B), confirming that they produced a functional gene product with no dominant negative effect through mixed oligomerization with WT PilB. In a strain lacking both *pilB* and *fimX*, all *pilB* Δ1bp truncations tested significantly increased twitching motility relative to the control complemented with WT PilB (Fig 3B). Notably, longer extensions rescued motility to a greater degree than shorter ones.

**Figure 4:**
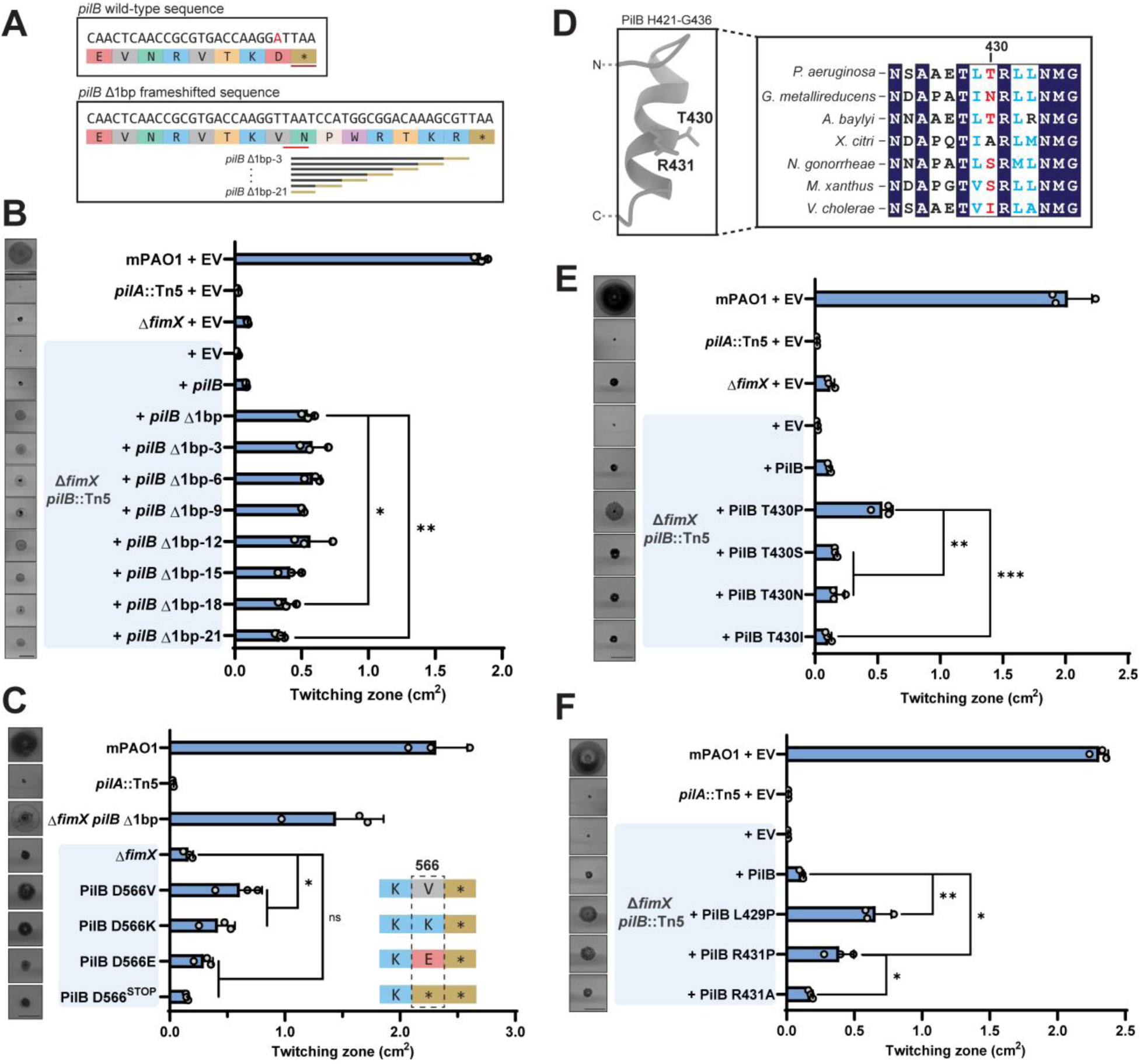
PilB mutations increase twitching motility in the absence of *fimX*. **(A)** Cartoon schematic of deleted base pair A1697 at the PilB C-terminus, shown in red, and resulting extended sequence. The *pilB* Δ1bp truncation sequences are shown below. **(B)** Quantification of twitching zone areas of Δ*fimX pilB*::Tn5 double mutants overexpressing *pilB* Δ1bp 3ʹ truncations. Representative crystal violet stained twitching zones are shown to the left. PilB expression was induced with 0.05% arabinose. Bars represent the means of duplicate samples from three independent experiments ± SD. **(C)** Quantification of twitching zone areas of Δ*fimX* PilB D566X point mutants chromosomal knock-ins. Representative twitching zones are shown to the left. Bars represent the means of triplicate samples from three independent experiments ± SD. **(D)** Full-length multiple sequence alignment of PilB homologues from various species highlighting the sequence surrounding PAO1 PilB T430, labelled in red. The Alphafold3 model of the predicted secondary structure in this region is shown to the left. **(E)** Quantification of twitching zone areas of *fimX pilB* double mutants overexpressing PilB T430X mutants. Representative twitching zones are shown to the left. PilB expression was induced with 0.05% arabinose. Bars represent the means of duplicate samples from three independent experiments ± SD. **(F)** Quantification of twitching zone areas of *fimX pilB*::Tn5 double mutants overexpressing PilB constructs with point mutations in positions adjacent to T430. Representative twitching zones are shown to the left. PilB expression was induced with 0.05% arabinose. Bars represent the means of triplicate samples from three independent experiments ± SD. **EV**: empty pHERD30T vector, ***pilB* Δ1bp-3 to 21**: *pilB* Δ1bp-3 to 21 in pHERD30T vector, **PilB L429/T430/R431X**: indicated PilB substitution in pHERD30T vector. All scale bars = 1 cm. **ns**: *p* ≥ 0.05; **\***: 0.05 ≥ *p* ≥ 0.01; **\*\***: 0.01 ≥ *p* ≥ 0.001; **\*\*\***: 0.001 ≥ *p* (Two-tailed parametric *t*-test).

Interestingly, the shortest *pilB* Δ1bp-21 truncation was identical in length to WT PilB, yet only the *pilB* Δ1bp-21 sequence significantly increased twitching motility in the double mutant (Fig 4B). Only the C-terminal amino acid differs between the two, with a valine substituted for an aspartate in the *pilB* Δ1bp-21 sequence. PilB D566 may therefore play an important role in modulating the function of the enzyme. The longer *pilB* Δ1bp sequences that support increased motility have additional positively charged residues (Fig 4A), suggesting that loss of negative charge at the C-terminus enhances PilB function. Since the above studies were done with multicopy plasmids, we next introduced single chromosomal point mutants at the PilB C-terminus in a Δ*fimX* background to assess recovery of twitching motility. A PilB D566V mutation significantly increased twitching motility compared to the Δ*fimX* parent (Fig 4C). We also observed a significant increase in twitching when the C-terminal amino acid was charge-swapped to a lysine, but not when we introduced glutamate (Fig 4C). The second-last amino acid in the native PilB sequence is a positively charged lysine, and we hypothesized that shortening the WT sequence by a single amino acid to expose the positive charge would also restore motility in a Δ*fimX* mutant. However, substituting D566 with a stop codon did not significantly increase motility relative to the control (Fig 4C). These data suggest that PilB C-terminal charge, in addition to C-terminal length, is an important regulator of PilB function.

### Determinants for PilB T430P function

We next examined a second PilB suppressor (PilB T430P) which mapped to a site distal to the C-terminus (Fig S8). A multiple protein sequence alignment of full-length PilB homologues from various species revealed that the region surrounding T430 was well conserved, although T430 itself was not (Fig 4D). We therefore reasoned that this amino acid was less important to overall PilB function or potentially a species-specific regulatory site. We made individual substitutions at T430, choosing amino acids found at this position in the other homologues. All mutant constructs complemented twitching motility in a *pilB*::Tn5 background and the WT (Fig S10). In a *pilB fimX* double mutant, only the original T430P suppressor supported twitching motility (Fig 4E). Therefore, disruption of local PilB secondary structure, rather than specific side-chain chemistry, is likely responsible for increased motility.

An Alphafold3 model of a PilB monomer predicted that residue T430 is located in the second turn of a short α-helix (Fig 4D and S8). While T430 is not well conserved, other amino acids in this helical segment, such as an adjacent R431, are. Alignment of the model onto the crystal structure of hexameric PilB from *Geobacter metallireducens* bound to a non-hydrolysable ATP analogue shows that the equivalent N429 (T430) and R430 (R431) residues are positioned at the protomer-protomer interface or jutting into the lumen of the torus, depending on the phase of the ATP hydrolysis cycle (Fig S8). We hypothesized that this secondary structural element could be involved in inter-subunit communication and that the introduction of a helix-breaking proline would alter these functional contacts. Supporting this idea, mutation of the adjacent residues to proline, L429P or R431P, significantly increased twitching motility in *fimX pilB* (Fig 4F). In contrast, R431A failed to increase motility in the double mutant, thus disruption of the α-helical segment rather than loss of this sidechain chemistry was likely responsible for altering PilB function. Interestingly, in the *pilB*::Tn5 or WT backgrounds where FimX was present (Fig S11A and B), twitching motility of PilB R431A and R431P variants was significantly decreased compared to control (Fig S11A). Therefore, the presence or absence of FimX modulates the function of these PilB variants.

### Increased twitching of suppressors is due to enhanced PilB ATPase activity

We next investigated how the *pilB* Δ1bp and PilB T430P mutations restored pilus function in a Δ*fimX* background. Other PilB regulatory effectors, such as PilZ, are thought to serve as molecular chaperones and *pilZ* mutants lack detectable intracellular PilB on Western blots (Fig 5A). This is consistent with results from previous studies of the *Xanthomonas citri* PilZ homologue (27). The *pilZ* mutants are also non-motile and phage resistant (Fig 1B and C), in keeping with their negligible levels of PilB. In contrast, PilB was still detectable in Δ*fimX* or FimX AAA backgrounds, although its levels were approximately half of WT (Fig 5A). Neither suppressor increased levels of PilB compared to those in Δ*fimX* or FimX AAA. Since FimX is important for PilB polar localization (8, 27, 30), the reduction in PilB levels in *fimX* mutants may be a consequence of fewer interactions between the ATPase and T4P machinery, making it more susceptible to degradation.

**Figure 5:**
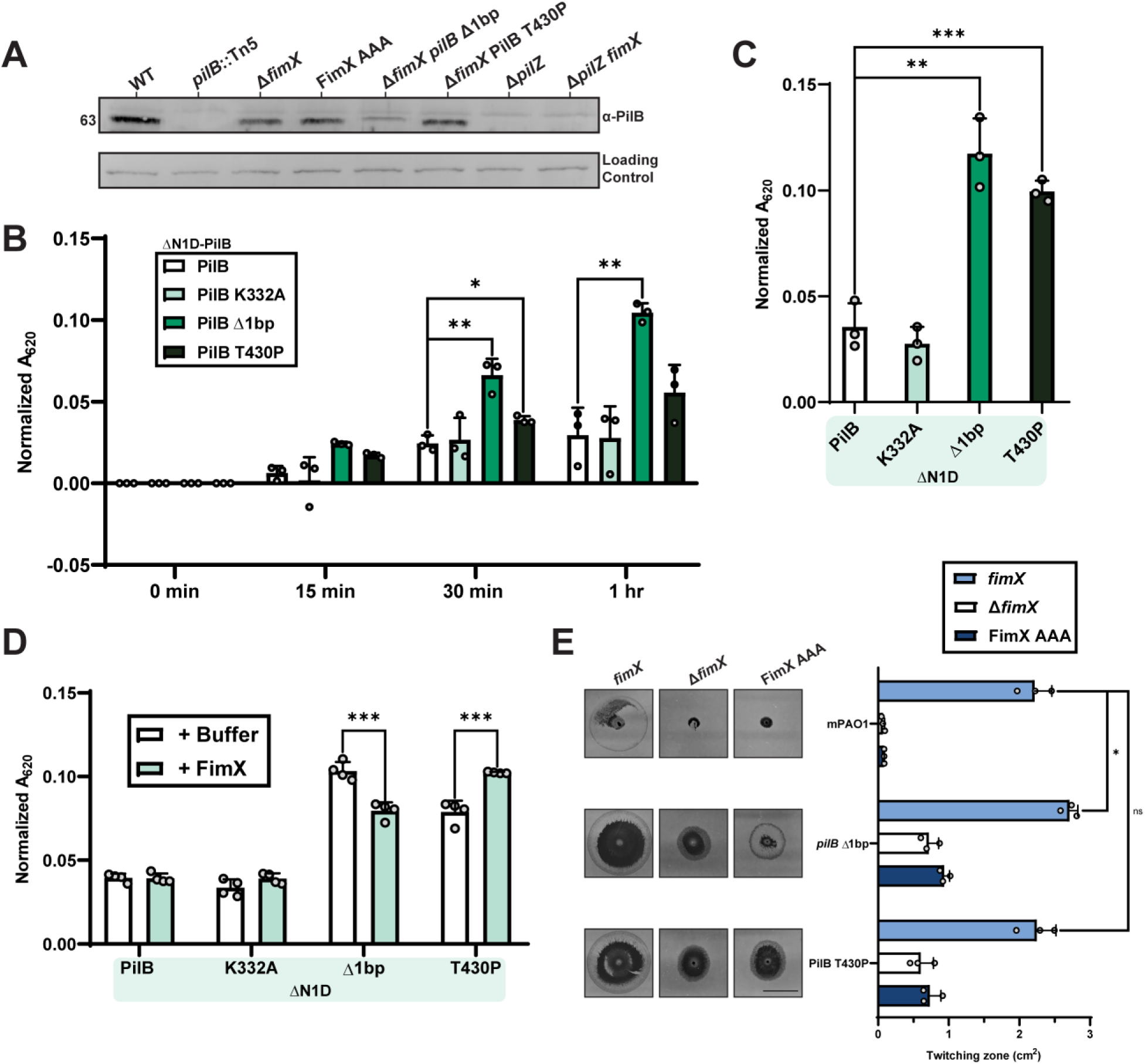
PilB suppressor mutations increase ATP hydrolysis. **(A)** Representative PilB Western blot. Sample loading was normalized to a non-specific band shown below. Molecular weight is indicated to the left. Blot is representative of three independent experiments. Quantification using BioMol green reagent of free phosphate released over a time course **(B)** or following overnight incubation **(C)** by ΔN1D-His_6_-PilB-catalyzed ATP hydrolysis. PilB K332A, a non-catalytically active Walker mutant, was used as a negative control. **(D)** Quantification of free phosphate release by ΔN1D- His_6_-PilB-catalyzed ATP hydrolysis in the presence of cdGMP containing reaction buffer (+Buffer) or with cdGMP and FimX (+FimX) following overnight incubation. **(E)** Quantification of twitching zone areas of Δ*fimX* or FimX AAA mutants with or without individual PilB suppressor mutation chromosomal knock-ins. Representative crystal violet stained twitching zones are shown to the left. Bars represent the means of triplicate samples from three independent experiments ± SD. Scale bar = 1 cm. **ns**: *p* ≥ 0.05; **\***: 0.05 ≥ *p* ≥ 0.01; **\*\***: 0.01 ≥ *p* ≥ 0.001; **\*\*\***: 0.001 ≥ *p* (Two-tailed parametric *t*-test).

Since PilB is an ATPase used to power pilus extension, we reasoned that the mutations might increase enzymatic activity. PilB has two N-terminal domains (N1D and N2D) connected by a flexible linker to the C-terminal domain (Figure S8) (43). The N1D can be removed without disrupting hexamer formation to improve stability and solubility for *in vitro* studies (27, 44–48). We purified ΔN1D-PilB variants to assess ATP hydrolysis efficiency *in vitro* (Fig S12). Both PilB suppressors hydrolyzed significantly more ATP than the WT and catalytically inactive Walker A mutant (PilB K332A) (1) controls over time (Fig 5B), as well as following overnight incubation (Fig 5C). Despite mapping to disparate sites in the protein, both PilB suppressors upregulated enzymatic activity and likely compensate for loss of FimX through this shared mechanism. Interestingly, when we added FimX and cdGMP to the reaction, we saw a decrease in ATP hydrolysis for *pilB* Δ1bp but an increase for PilB T430P (Fig 5D). While this result is difficult to interpret, it suggests that FimX can modulate PilB enzymatic activity. In a genetic context, *pilB* Δ1bp and PilB T430P mutations integrated into the chromosome restored twitching motility to at least WT levels in the presence of FimX (Fig 5E), but not FimX AAA. Taken together, this indicates that these PilB suppressor mutants can continue to respond to cdGMP- bound FimX input and regulation.

### FimX promotes pilus extension to produce longer pili

To test the impact of increased ATP hydrolysis by PilB on pilus assembly, we imaged pilus dynamics using fluorescence microscopy (Fig 6A). The major pilin subunit, PilA, was first labelled with a fluorescent maleimide dye at A86C as previously described (4, 49). All PilA A86C mutants remained motile with slightly reduced twitching compared to the wild type (Fig S13). WT cells produced T4P at the cell poles as expected, with some cells producing pili at both poles (Fig 6A). Notably, the Δ*fimX* mutant produced very short pili that were difficult to detect, often presenting only as bright polar puncta which we scored as a polar accumulation of labelled PilA or very short T4P, below the limit of detection. In addition, fewer cells were piliated compared to WT (Fig 6B and C). We chose to quantify pili using the Δ*fimX pilB* Δ1bp mutant as this strain showed the most robust twitching phenotype of our suppressors. Furthermore, in the presence of FimX, the *pilB* Δ1bp mutation alone had greater than WT levels of twitching, providing an optimal background to study FimX’s functional contribution. In the Δ*fimX pilB* Δ1bp background, piliation was restored on many cells (Fig 6B and C), with longer pili compared to the Δ*fimX* mutant (Fig 6A). The percentage of piliated cells in the Δ*fimX pilB* Δ1bp mutant was similar to WT (Fig 6B and C), but the average maximum pilus length was significantly shorter (Fig 6D), likely explaining its comparatively reduced motility (Fig 5E). Maximum pilus length was significantly increased compared to the Δ*fimX* parent, indicating that the *pilB* Δ1bp mutation caused cells to produce longer pili, presumably through enhanced ATP hydrolysis. The *pilB* Δ1bp alone background produced the longest pili (Fig 6A and D) and had the highest percentage of piliated cells (Fig 6B and C), consistent with this strain being the most motile (Fig 5E). Longer and long-lived pili in PAO1 (∼2-3 µm) comprise a rare subpopulation (4). We therefore anticipate that more Δ*fimX* cells are likely piliated, but since the maximum pilus length in these mutants is ∼0.75 µm, most pili may not be detectable.

**Figure 6:**
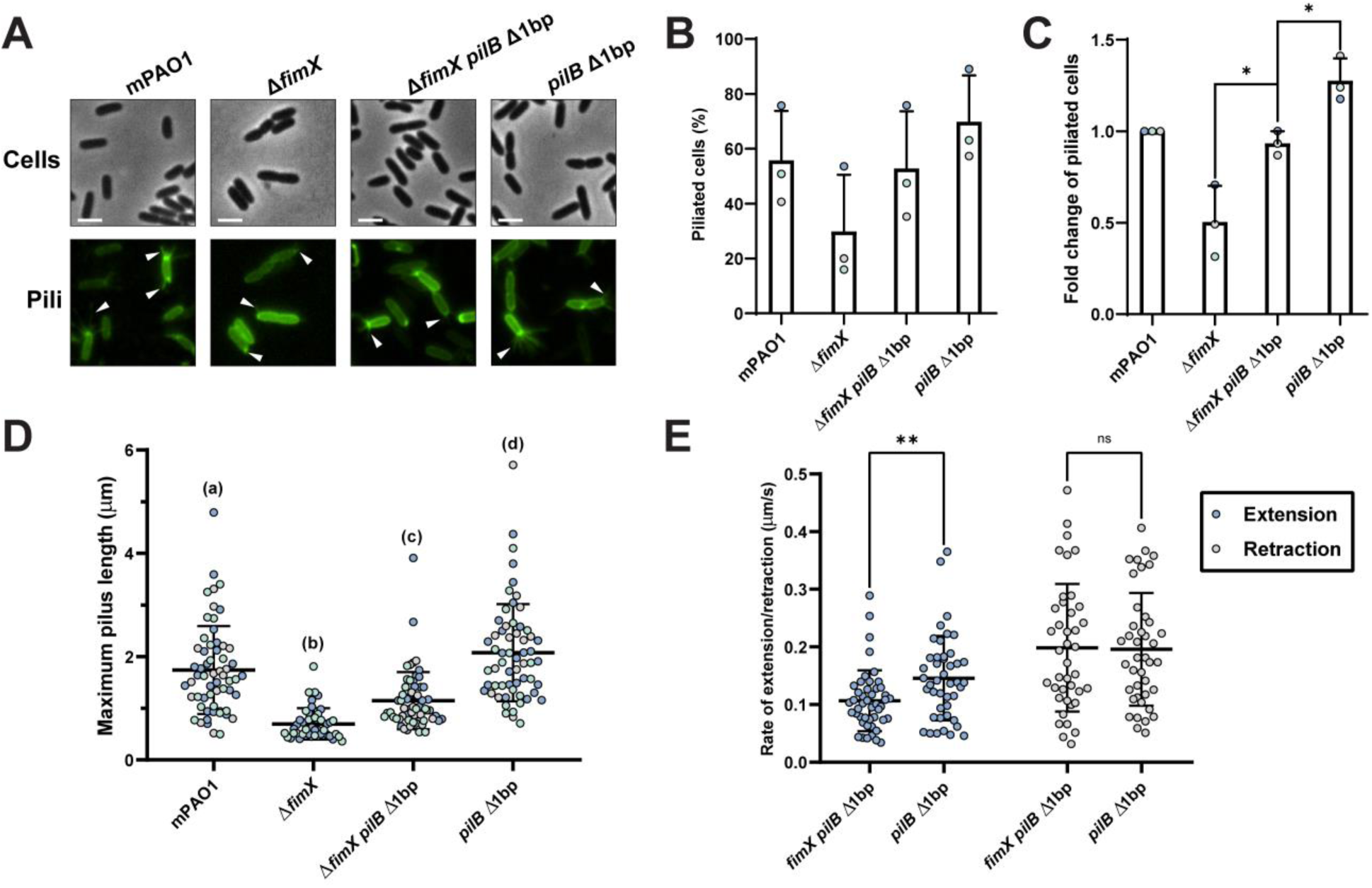
FimX mutants produce short pili due to slow extension rates. **(A)** Representative images of PilA A86C type IV pili labelled with maleimide-Alexafluor 488 with background fluorescence subtracted. White arrows indicate visible pili. All scale bars = 2 µm. Number of cells producing visible PilA A86C type IV pili expressed as a percentage of total cells **(B)** or normalized relative to the WT arbitrarily set to 1 **(C)**. Bars represent the means ± SD of three independent experiments (WT *n* = 193, Δ*fimX n* = 181, Δ*fimX pilB* Δ1bp *n =* 210, *pilB* Δ1bp *n =* 197 total cells). Technical replicates are indicated by colour. **(D)** Maximum PilA A86C pilus length. Each sample depicts the means ± SD of three independent experiments, each point represents a single PilA A86C pilus length (WT, Δ*fimX pilB* Δ1bp, and *pilB* Δ1bp *n* = 60, Δ*fimX n* = 45 total pili). Technical replicates are indicated by colour. **(E)** Rate of PilA A86C pilus extension or retraction. Each sample depicts the means ± SD of three independent experiments, each point represents the rate of extension and/or retraction of a single PilA A86C pilus (Δ*fimX pilB* Δ1bp *n* = 46, *pilB* Δ1bp *n* = 44 total extension and Δ*fimX pilB* Δ1bp *n =* 40, *pilB* Δ1bp *n =* 41 total retraction events). **ns**: *p* ≥ 0.05; **\***: 0.05 ≥ *p* ≥ 0.01; **\*\***: 0.01 ≥ *p* ≥ 0.001 (Two-tailed parametric *t*-test).

The *pilB* Δ1bp mutants afforded an opportunity to quantify pilus dynamics in the presence or absence of FimX. We quantified the rates of pilus extension and retraction over time in the Δ*fimX pilB* Δ1bp versus *pilB* Δ1bp to assess FimX’s contribution to pilus dynamics. The FimX-expressing strain had a significantly increased average rate of pilus extension (Fig 6E) while the average rate of retraction was not significantly different, supporting FimX’s role as a modulator of PilB-dependent pilus extension. These data suggest that FimX directly increases pilus extension efficiency, likely by acting on PilB.

In *P. aeruginosa* and other bacterial species, fluorescently labelled FimX fusions display defects in polar localization when PilB interaction is impaired (27, 30, 33). Consistent with these other studies, most Δ*fimX* mutant cells complemented with an N-terminal fusion of mNeonGreen (mNGr) to FimX showed bright puncta localized to a single cell pole (Fig 7A), with a smaller population of cells displaying bipolarly-distributed puncta (Fig 7B). The WT FimX fusion also complemented twitching motility (Fig S14A), while cells expressing an empty vector control had no discernable puncta (Fig S14B). The number of cells with identifiable polar puncta was decreased by ∼20-50% when *pilB* was deleted or if a mNGr-FimX AAA fusion was used (Fig 7B). Importantly, these data indicate that FimX localization patterns in our strains are consistent with previous reports (8, 27, 30) and are contingent on interaction with PilB.

**Figure 7:**
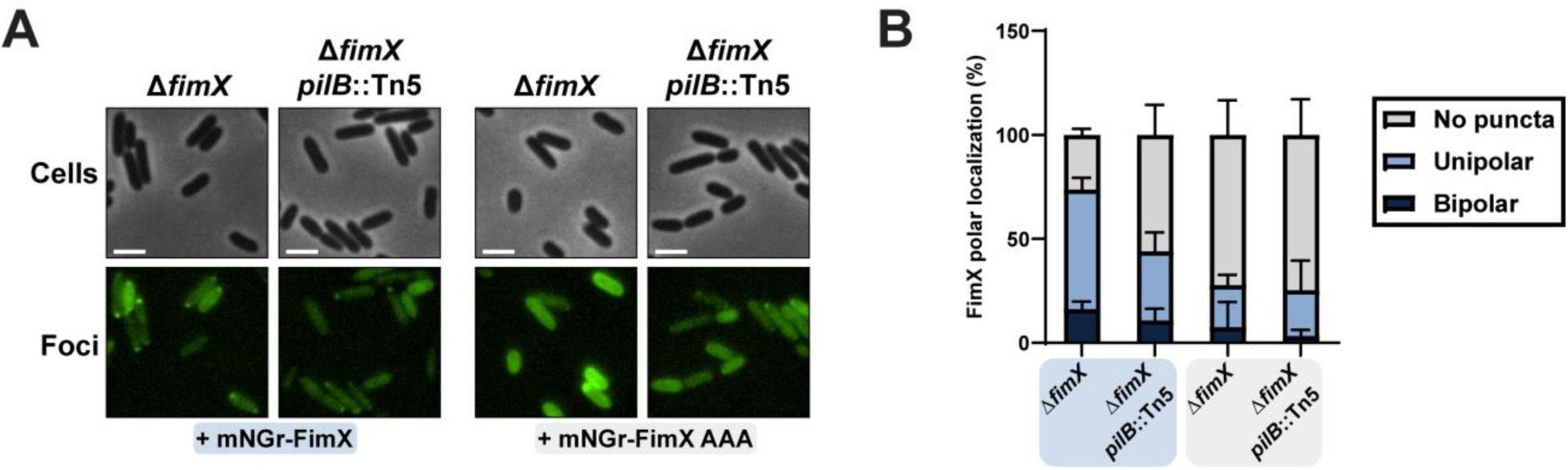
FimX polar localization is disrupted in the absence of PilB binding. **(A)** Representative images of cells expressing an mNeonGreen N-terminal FimX or FimX AAA fusion with background fluorescence subtracted. All scale bars = 2 µm. **(B)** Quantification of cells expressing an mNeonGreen-N-terminal FimX or FimX AAA fusion scored for bipolar, unipolar, or no discernable puncta and expressed as a percentage of total cells. Bars represent the means ± SD of three independent experiments (Δ*fimX* + mNGr-FimX *n=* 216, Δ*fimX pilB*::Tn5 + mNGr-FimX *n=* 213, Δ*fimX* + mNGr-FimX AAA *n=* 235, Δ*fimX pilB*::Tn5 + mNGr-FimX AAA *n=* 206, total cells). **mNGr-FimX** or **mNGr-FimX AAA**: mNeonGreen N-terminal FimX or FimX AAA fusion in pHERD30T vector. Expression was not induced with arabinose.

### PilB enhancing mutations are uncommon

Given the relative ease of selection for *fimX* suppressor mutations that mapped to PilB and improved twitching motility, we wondered if similar PilB mutations were common in sequenced strains of *P. aeruginosa*. We compared 417 PilB orthologue sequences from the Pseudomonas.com database to identify comparable mutations. Of these, 297 were the same length as PAO1 PilB at 566 residues, and 82 were longer. Of those sequences, 72 were longer by one residue, 8 by two residues, 1 by four residues, and 1 by 15 residues (Fig S15A). Strikingly, despite these differences in length, the C-terminal residue of all sequences was an aspartic acid. We also assessed the local sequence surrounding PilB T430 for helix-disrupting residues which may confer a similar effect to T430P. We examined the region +/- three amino acids from the residue aligned with PAO1 PilB T430 in a full-length protein sequence alignment, to capture the most likely positions to influence stability of a 3-turn α-helix. Of the 417 sequences, 4 had a proline residue and 8 had a glycine within this window, accounting for less than 1% and 2% of the total sequences respectively (Fig S15B). This analysis of *P. aeruginosa* sequences suggests that PilB mutations similar to the ones identified in this screen are uncommon.

## Discussion

The reason that optimal PilB function in *P. aeruginosa* needs the regulatory input of FimX remains poorly understood (30, 31). To address this knowledge gap, we screened for suppressors capable of overcoming the Δ*fimX* motility deficit. We identified several suppressor mutants, most of which mapped to the Pil-Chp network or to PilB (Table S1). A previous Tn-Seq screen used to isolate suppressors in *P. aeruginosa* strain PA103 implicated FimX as a sensor of the secondary messenger cdGMP. In that strain, Δ*fimX* suppressors with high levels of cdGMP had increased motility, leading the authors to posit that FimX senses internal cdGMP to tune pilus assembly efficiency over broad concentrations of the molecule (31). Unexpectedly, we found no suppressors in cdGMP homeostatic machinery, nor did any of our mutations increase cdGMP synthesis (Fig 2A). Further, exogenous expression of DGCs also failed to alleviate the Δ*fimX* motility defect in PAO1 (Fig 2B), unlike in PA103 Δ*fimX* where expression of the unrelated DGC PleD from *Caulobacter crescentus* was sufficient to restore motility and surface piliation. While these phenotypic discrepancies may be strain-specific, overexpression of other DGCs in PAO1 was reported to suppress twitching motility, though surface piliation was not significantly impacted (31). Why these strains behave differently in response to cdGMP production is not yet clear.

Instead, we found that increases in the secondary messenger cAMP increased twitching in Δ*fimX*, though this is likely an indirect mechanism. Increasing the levels of cAMP drives transcription of the Vfr regulon, which includes T4P components. Mutations that promote accumulation of cAMP lead to the expression of additional pilus machines. Given that Δ*fimX* mutants retain the ability to twitch, albeit poorly, these high cAMP mutants likely produce an abundance of sub-optimally functioning T4P machines which cumulatively improve the cells’ ability to twitch (Fig 8). A similar pattern of compensation via elevated cAMP production was recently reported for suppressors of another modulator of pilus function in strain PA14 (50).

**Figure 8:**
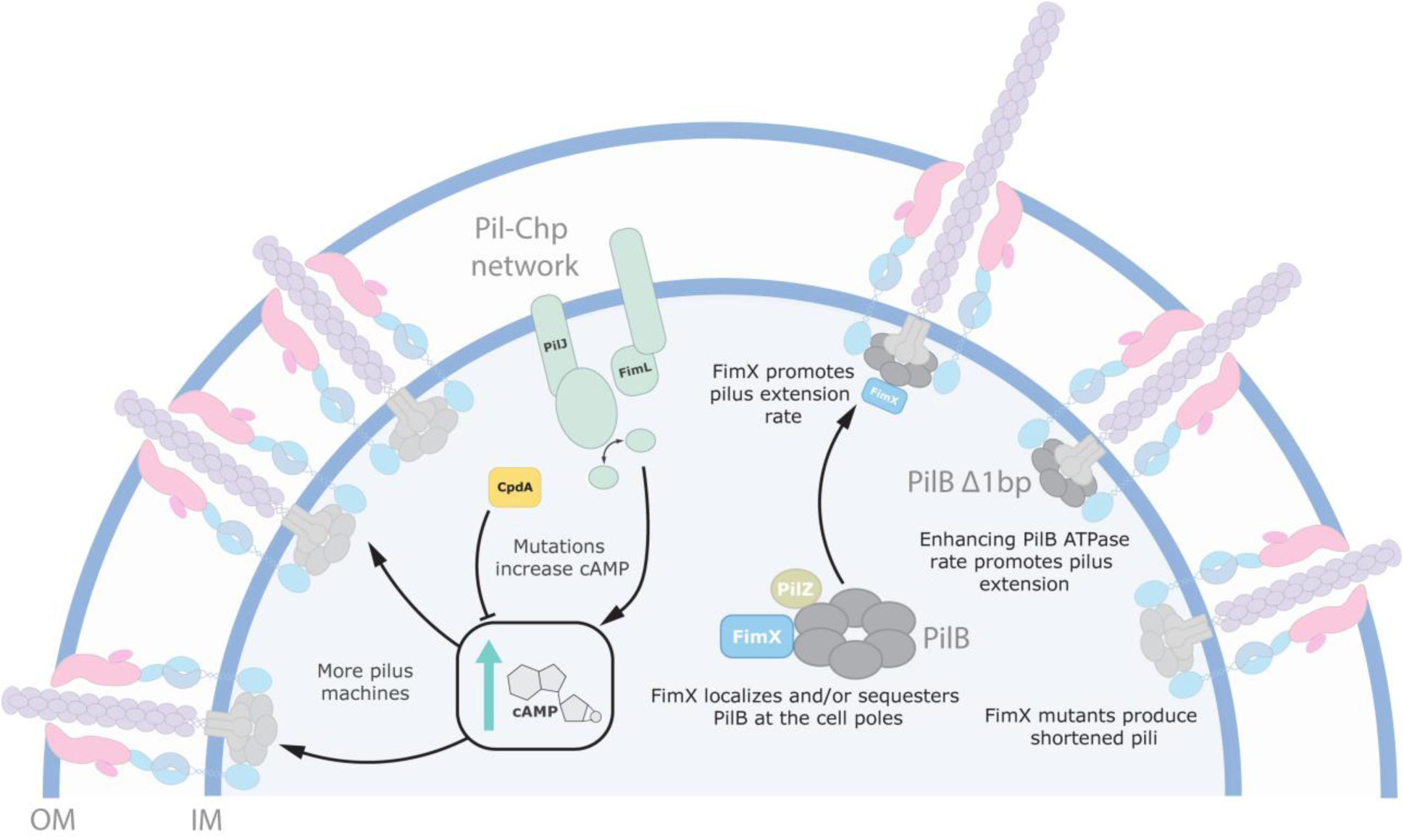
Model of Δ*fimX* suppression pathways. Suppression of the Δ*fimX* twitching motility deficit can occur through mutations which elevate intracellular levels of the secondary messenger molecule cAMP. More cAMP generation leads to increased Vfr-dependent expression of T4P components, which assemble into more machines, increasing twitching motility. This mechanism is not FimX-specific. Alternatively, mutations that increase PilB ATP hydrolysis lead to production of longer pili. This overcomes the short-pilus defect observed in *fimX* mutants. FimX in complex with PilB enhances the rate of PilB-dependent pilus extension to produce longer pili optimized for twitching motility. FimX polar localization to T4P machines is contingent on PilB binding, therefore FimX influences its polar trafficking and/or sequestration.

We also isolated a suppressor that mapped to PilO (PilO P168L), a member of the T4P alignment subcomplex (35). Notably, this mutant was the only one tested that was not complemented to at least WT twitching by FimX expression (Fig S4). Therefore, the twitching recovery mechanism for this mutant is likely independent of the FimX-PilB extension pathway. PilO interacts with regulatory components such as SadC (18) and may connect pilus function to environmental signals.

The suppressor mutations in PilB provided a useful tool to study FimX-dependent physiology. Despite mapping to separate sites (Fig S8), both mutations increased ATP hydrolysis (Fig 5A and B) suggesting that one of FimX’s roles may be to enhance PilB enzymatic activity. For the *pilB* Δ1bp mutant, mechanistic clues come from structural work on other extension ATPases. The C-terminal solvent-exposed face of the *G. metallireducens* PilB hexamer has dispersed negative surface charge (25, 26). The extended Δ1bp C-termini, which include several positively charged residues, are predicted to protrude into this environment (Fig S8). Increasing positive charge, or removing the native C-terminal negative charge, could promote electrostatic stabilization of the hexamer, or facilitate the PilB conformational changes necessary for substrate hydrolysis.

Analysis of the PilB T430P mutant uncovered a novel regulatory region in the PilB multimer. Disruption of the short helix containing T430 restored twitching motility in a Δ*fimX* background. The adjacent R431 residue is highly conserved in PilB homologues, and an R431P variant increased twitching in Δ*fimX* but interestingly, significantly attenuated motility in the presence of FimX (Fig S11). These PilB mutants therefore continue to interact with and respond to FimX, though resulting in different functional outcomes. This secondary structural element may represent an intramolecular regulatory site involved in FimX recognition or perhaps is involved in communicating conformational changes induced through FimX interaction. More detailed biochemical studies are still required to unravel these possibilities.

In the absence of FimX, twitching is diminished. Using fluorescence microscopy, we showed that the pili of *fimX* mutants are significantly shorter than WT (Fig 6D). This result is consistent with low levels of surface pili in sheared protein fractions (Fig S1). The Δ*fimX pilB* Δ1bp suppressor mutant had increased PilB ATP hydrolysis and produced longer pili, presumably through increased PilA polymerization. These data suggest that twitching motility in PAO1 is maximized when cells make longer pili, as WT and *pilB* Δ1bp strains generated the longest pili and largest twitching zones. When comparing Δ*fimX pilB* Δ1bp and *pilB* Δ1bp, we saw a significant increase in the rate of pilus extension, but not retraction (Fig 6E). Our data therefore support a model where FimX-PilB interaction stimulates pilus extension, potentially by stabilizing or promoting the PilB conformational changes necessary to drive pilus assembly within the stator (Fig 8). In *X. citri*, the addition of FimX stimulated ATP hydrolysis by a PilB-PilZ complex *in vitro* (27). We also showed that FimX influenced *P. aeruginosa* ΔN1D-PilB function *in vitro* (Fig 5D), implying that the two proteins interact. In other bacteria, FimX is thought to engage the extension machinery through PilZ (27, 33, 51, 52). Similar FimX-PilZ interactions have not been detected in *P. aeruginosa* (53), however, leaving definition of the specific interaction interfaces for future studies.

In *P. aeruginosa* and *Xanthomonas spp*., FimX has also been proposed to function as a PilB polar localization factor (8, 27, 30), rather than as a chaperone like PilZ. In *P. aeruginosa*, FimX shows polar localization defects in *pilB* backgrounds or when it is unable to bind to cdGMP (Fig 7), suggesting that both cdGMP-bound FimX and PilB are required for polar co-localization (30). The mechanism behind polar trafficking is poorly understood, but recent evidence suggested that the Pil-Chp response regulator PilG was essential for proper PilB polar localization (8, 34). Consistent with these observations, a recent study in *Acinetobacter baylyi* demonstrated that its FimX homologue controls T4P localization through interactions with components of the Pil-Chp pathway, but that this localization is independent of cdGMP binding (32). How those factors are targeted to the cell poles in *P. aeruginosa* remains an open question. PilB and FimX may first need to form a complex which is then recognized by additional factors such as PilG. Alternatively, FimX interaction may bias PilB polar occupancy or keep it sequestered, ensuring that the leading pole is primed for multiple rounds of pilus assembly (Fig 8). In this scenario, PilB and FimX polar retention is mutually contingent, but FimX instead functions to reduce disengagement of PilB from active T4P machinery.

In other bacteria such as *Vibrio* or *Neisseria* species, pili are used predominantly for surface adhesion (54, 55), cell-cell contact (56), DNA uptake (57, 58), or biofilm formation (55). These organisms lack obvious FimX homologues. Instead, the N-terminal domain of extension ATPases in these species binds directly to cdGMP, and nucleotide binding is essential for function (54, 59–61). Pilus extension is therefore optimal at high intracellular cdGMP concentrations, but this condition suppresses twitching motility. In *P. aeruginosa,* nucleotide binding to FimX, which then dictates PilB localization and function, may serve as a functional alternative. As a high affinity sensor of cdGMP (29, 33), FimX may drive optimal pilus extension rates to enable multiple dynamic cycles of pilus assembly, and therefore facilitate twitching, in response to nucleotide availability. This organization might also explain why mutations which enhance PilB function are rare in *P. aeruginosa*, as these may dysregulate pilus functional responses to signaling cues.

## Materials and Methods

### Bacterial strains and growth conditions

All strains and plasmids used in this study are outlined in Table S2. Bacterial strains were grown aerobically at 37 °C in Lysogeny Broth (LB) media (10 g/L tryptone, 5 g/L yeast extract, and 5 g/L NaCl), supplemented with the appropriate concentration of antibiotics where necessary unless otherwise specified. Antibiotics were used at the following concentrations: 15 µg/mL or 30 µg/mL of gentamicin for *E. coli* or *P. aeruginosa* respectively unless otherwise indicated. 50 µg/mL or 200 µg/mL kanamycin for *E. coli* or *P. aeruginosa* respectively. 100 µg/mL of ampicillin or 200 µg/mL of carbenicillin for *E. coli* or *P. aeruginosa* respectively. Plasmid constructs were generated using *E. coli* DH5α before introduction into the appropriate cellular background by heat shock (*E. coli*) or electroporation (*P. aeruginosa* or *E. coli*) where appropriate.

### Plasmid and strain construction

Fragments of interest were PCR amplified from isolated mPAO1 genomic DNA. All PCR primers are listed in Table S3. For point mutations, two separate PCR amplified DNA fragments were stitched together through a second single overlap extension PCR, with each DNA overhang including the mutation of interest introduced during primer synthesis. For plasmid construction, the in-frame genetic insert and corresponding vector were digested with the appropriate FastDigest restriction enzymes (ThermoFisher Scientific) as per manufacturer instruction and ligated with T4 DNA ligase. Ligated products were transformed via heat shock into *E. coli* DH5α and selected on 1.5% agar LB media plates supplemented with the appropriate antibiotic. Following selection, single colonies were harvested, and the plasmid was mini prepped using a GeneJet plasmid prep kit (ThermoFisher Scientific) for sequence confirmation (Plasmidsaurus).

Genetic manipulation of mPAO1 chromosomal DNA was performed by double homologous recombination as previously described (62). In brief, for genetic deletions, ∼500 bp upstream and downstream of the desired target sequence were PCR amplified and purified from isolated PAO1 genomic DNA. Upstream and downstream sequences were engineered to contain ∼50-100 bases of the 5ʹ or 3ʹ ends of the gene deletion target to ensure minimal disruption of adjacent genes. Upstream and downstream fragments were digested with the appropriate restriction enzymes along with the suicide vector backbone pEX18Gm. Digested fragments were ligated using T4 ligase and transformed into via heat shock into *E. coli* DH5α and selected for on 1.5% agar LB media plates supplemented with gentamicin. Purified and sequenced plasmids were then transformed into *E. coli* SM10 (λ*pir*). A single colony of SM10 harbouring the plasmid was grown overnight in 3 mL of LB media supplemented with gentamicin. The following day, the SM10 culture was mixed with the appropriate strain of mPAO1 overnight culture in a 1:1 ratio (by volume), pelleted by centrifugation, the resulting cell pellet was resuspended in 50 µL of LB and then spotted on 1.5% agar LB plates dried in a biological safety cabinet for 20-30 min. The spot was allowed to dry at room temperature before being incubated at 37 °C overnight. The following day, the cell spot was scraped into fresh LB media and plated onto premixed *Pseudomonas* isolation agar (Difco) supplemented with 100 µg/mL of gentamicin. The plates were incubated again at 37 °C overnight. Single colonies were then re-streaked again on *Pseudomonas* isolation agar plates supplemented with 100 µg/mL of gentamicin and incubated overnight at 37 °C. The next day, cells were streaked in a zig-zag pattern on 10% LB (1 g/L tryptone, 0.5 g/L yeast extract) with no added NaCl, 1.5% agar, and 5% sucrose media plates and incubated overnight at 37 °C. Single colonies from the sucrose media selection plate were patched again onto a fresh sucrose media plate as well as an LB media plate supplemented with gentamicin. Patches which grew on sucrose media but not LB with gentamicin were assessed for chromosomal deletion via colony PCR.

For chromosomal knock-in of a point mutation, the same protocol as outlined above was performed only differing at initial construct design. Knock-in constructs were instead generated ∼500 bp upstream or downstream of the mutation site through single overlap extension PCR. Overlap extension primers were also designed to silently introduce or remove a restriction cut site in the genome where applicable to facilitate downstream quality assurance. The remaining procedure was then carried out as described.

### Twitching motility assays

Bacterial strains were streaked onto 1.5% agar LB media plates supplemented with gentamicin where applicable and grown overnight at 37 °C. The following day, cells were picked using a sterile pipette tip and stabbed down to the bottom layer plastic of 1% agar LB media tissue culture treated plates, supplemented with antibiotic or arabinose where applicable, dried for 30 min in a biological safety cabinet. Plates were incubated overnight at 37 °C. The following day, the agar was carefully removed with a pipette tip and discarded. The twitching zones adhered to the plate were stained with 0.1% crystal violet dye for a minimum of 10 min, washed twice with DI water, and left to air dry overnight at room temperature before imaging with a scanner. Twitching zone areas were then digitally traced using ImageJ, and either duplicates or triplicates were averaged.

### Phage-susceptibility assays

Bacterial strains were inoculated into 3 mL of LB media liquid cultures and grown overnight at 37 °C. The next morning, 5 µL of overnight culture was sub-cultured into 3 mL of fresh LB media and grown at 37 °C to mid-logarithmic phase (OD_600_ ∼0.5-1.0). Cells were normalized to an OD_600_ of 0.1 in LB. For plate-based assays, 10 µL of bacterial suspension was mixed with 5 µL of serially diluted PO4 phage stock (3e5 PFU/mL original titer). 10 µL of cell-serial phage dilution was spotted onto 1.5% agar LB media dried for 30 min in a biological safety cabinet. The plate was incubated overnight at room temperature. The following day, the plate was incubated again at 37 °C for 2-3 hours before imaging with a scanner.

For liquid-based assays, 2 µL of OD_600_ normalized cell suspension was added to 200 µL of LB media total in a flat bottom 96 well plate. To each well, 1 µL of 3e5 PFU/mL PO4 phage stock was also added. The plate lid was sealed with masking tape and then incubated overnight in a 37 °C shaking plate reader, with OD_600_ readings taken every 30 min for 16-18 hours. Curves were normalized to LB media blank wells in the same plate.

### Sheared surface protein preparation

Surface pilin and flagellin protein levels were assessed as previously described (63). Briefly, strains were streaked in grid-like pattern on a 1.5% agar LB media plate dried for 30 minutes in a biological safety cabinet. The plate was incubated at 37 °C overnight. The next morning, cells were gently scrapped off the surface using a combination of a glass coverslip and sterile inoculation loop into 1.2 mL of sterile 1X PBS. Surface appendages were sheared by vortexing the suspension for 30 s. The homogenized suspension was transferred to a 1.5 mL Eppendorf tube and pelleted by centrifugation at maximum speed for 30 min. The supernatant was decanted into a fresh tube and 0.1 volume of 5M NaCl and 30% (w/v) polyethylene glycol (MW range 8k) was added. The tubes were then incubated on ice for 90 min, inverting the tubes periodically. The precipitated protein was collected by centrifugation again for 30 min at maximum speed. The supernatant was decanted, and the precipitated protein resuspended in 50 µL of 1X SDS loading dye (62.5 mM Tris-HCl pH 6.8, 0.5% (w/v) SDS, 5% (v/v) β-mercaptoethanol, 10% (v/v) glycerol, 0.25% (w/v) bromophenol blue), and boiled for 10 min. The tubes were centrifuged for another 5 min at maximum speed and separated on a 15% SDS-polyacrylamide gel. Proteins were visualized with Coomassie blue staining.

### Bacterial 2-hybrid assay

Single colonies of *E. coli* BTH101 co-transformed with both T18/T25 tagged target proteins were inoculated into 3 mL LB media overnight cultures supplemented with kanamycin, ampicillin, and 0.5 mM IPTG. The following morning, the cultures were normalized to an OD_600_ of 0.1 in fresh LB and 4 µL was spotted onto MacConkey agar plates (40 g/L, Difco) supplemented with kanamycin, ampicillin, 0.5 mM IPTG, and 1% maltose pre-dried for 40 min in a biological safety cabinet. The spots were allowed to dry at room temperature before incubation at 30 °C in a standing incubator. The plates were imaged after 24 hours.

### Selecting Δ*fimX* twitching suppressor mutants

Motility suppressors were isolated through the standard twitching assay described with some alterations. Δ*fimX* bacteria were first streaked for single colonies on an LB agar plate. Using a sterile pipette tip, a single colony was stab-inoculated into a standard twitching plate ∼4-5 times in a row. The tip was discarded, and a new colony was then stab inoculated in the same manner. This was repeated for a total of 35-40 individual twitching zones on a single assay plate. The plate was sealed with parafilm and incubated at 37 °C for 48-72 hours. Following incubation, the top agar layer was carefully removed and discarded. Flares emerging from small twitching zones were collected with a sterile cotton swab and streaked again for single colonies. Single mutant colonies were reassessed for twitching motility. Those with increased twitching relative to the Δ*fimX* strain were collected and their genomic DNA isolated using a Wizard genomic DNA isolation kit (Promega) as per manufacturer’s instructions. Isolated chromosomal DNA was sent for Illumina Sequencing (SeqCenter) and mutations were identified using BreSeq (64). Where appropriate, candidate mutations were reintroduced into the chromosome as described above.

### cdGMP reporter assay

The cdGMP reporter assay was conducted as previously described (20) with some alterations. Briefly, cells transformed with the reporter construct (*pcdrA*:LuX-PMS402) were grown in LB media liquid cultures supplemented with kanamycin (as well as gentamicin for the high control carrying *sadC*-pBADGr) overnight at 37 °C. Overnight cultures were diluted (1:25) in 3 mL of fresh LB media with antibiotics and grown at 37 °C to mid-logarithmic phase. Cultures were then normalized to an OD_600_ of 0.1 and 2 µL of suspension was added to 200 µL of LB media total supplemented with antibiotics in a black flat clear-bottom 96 well plate. The plate was sealed with masking tape and then incubated overnight in a 37 °C shaking plate reader, with chemiluminescence and OD_600_ readings taken every 15 min for 16-18 hours. Relative luminescence signal was normalized to OD_600_ reading at the same timepoint.

### cAMP reporter assay

The cAMP reporter assay was conducted as previously described (37) with some alterations. Briefly, cells transformed with the reporter construct (*rpoD*:mKate/*PaQa*:YFP – O’Toole Lab) were grown in LB media liquid cultures supplemented with carbenicillin (as well as gentamicin for the high cAMP control carrying CyaB R456L-pBADGr) overnight at 37 °C. Overnights were sub-cultured 1:25 in 5 mL of M8 minimal media containing 0.2% glucose, 0.5% casamino acids, and 1 mM MgSO_4_ supplemented with carbenicillin to an OD_600_ of 0.2-0.4. At this point, 200 µL of liquid culture was spread onto 1% agar M8 media plates dried for 20 min in a biological safety cabinet. Liquid cultures and plates were then incubated for an additional 5 hours at 37 °C. After the incubation, 1 mL of sterile PBS was added to each plate and cells were scraped up with a sterile inoculation loop. 100 µL of cell suspension was taken from each plate and normalized to an OD_600_ of ∼0.2 in sterile 1X PBS. The liquid cultures were also normalized to an OD_600_ of ∼0.2 in sterile 1X PBS. 200 µL of normalized cell suspensions were then added to a black flat clear bottom 96 well plate (Corning). mKate (Ex: 588/Em: 635) and YFP (Ex: 500/Em: 541) fluorescence were measured, and the signal was normalized by dividing YFP/mKate for both surface-adapted and liquid samples.

### Vfr reporter assay

For *vfr* promoter activity detection, cells harbouring a gene encoding mRuby3 chromosomally integrated upstream of *pilM* and under control of the *pilMNOPQ* promoter were streaked onto 1.5% agar LB media plates and incubated overnight at 37 °C. The next morning, cells were harvested into sterile 1X PBS and normalized to an OD_600_ of ∼0.1. 200 µL of cell suspension was then added to a black flat clear bottom 96 well plate. mRuby3 (Ex: 558/Em: 600) fluorescence and exact OD_600_ were then measured, and the signal was normalized by dividing mRuby3/OD_600_.

### Western blot analysis

Whole cell lysates were first separated on a 15% or 12% SDS-polyacrylamide gels for PilA and PilB respectively. Samples were transferred to a nitrocellulose membrane at 225 mA for 1 hour. The membranes were blocked with 5% (w/v) skim milk (Bioshop) dissolved in 1X Tris-buffered saline (TBS: 20 mM Tris-HCl pH 7.6, 150 mM NaCl) overnight or for 2 hours for PilA or PilB respectively. Primary incubation was performed as previously described using rabbit antisera (65). The membranes were washed 4 times for 5 min with 1X TBS before incubation using alkaline phosphatase-conjugated goat-anti rabbit IgG secondary antibody (BioRad) in a 1:3500 dilution. The membranes were washed 4 times with 1X TBS and developed using alkaline phosphatase developing reagent according to manufacturer’s protocol, before imaging the membranes in an Azure Biosystems 400 transilluminator.

### Skim milk T2SS secretion assay

Liquid cultures were grown overnight in LB media at 37 °C. The next morning, each culture was normalized to an OD_600_ of 0.2 and spotted onto tryptic soy agar plates supplemented with 1.5% skim milk powder (Bioshop) dried for 30 min in a biological safety cabinet. The spots were allowed to dry at room temperature before incubation overnight at 37 °C. The resulting zones of clearance were then imaged with a scanner and the zone areas were quantified digitally using ImageJ.

### Protein modelling and sequence alignments

Alphafold3 was used to predict protein structures and generate predicted aligned error plots using the unmodified default settings (66). All models and plots were visualized in UCSF ChimeraX (67). Full length PilB sequences were aligned using Geneious Prime and visualized using ESPript3 (68).

### Protein purification

N-terminally truncated PilB (ΔN1D-PilB: M1-D180) and full length FimX were subcloned into pET28b with an N-terminal His_6_-tag for inducible expression. PilB was purified as previously described with some alterations. Briefly, ΔN1D-PilB expressing overnight cultures of BL21-CodonPlus (DE3)-RIL were sub-cultured (1:100) in 1 L of LB supplemented with kanamycin and incubated at 37 °C with shaking until mid-logarithmic phase (∼0.4-0.8 OD_600_). Cells were induced with 0.5 mM IPTG overnight at 18°C with shaking. The next morning, cells were harvested by centrifugation at 3000 xg for 20 min. Cell pellets were resuspended in lysis buffer (50 mM HEPES pH 7.3, 250 mM NaCl, 10 mM imidazole) and incubated on ice with 20mg of lysozyme for 15 min with periodic inversion. Cells were then lysed using sonication and the supernatant clarified through centrifugation at 31000 xg for 30 min. Supernatant was then incubated with ∼5 mL of Ni^2+^-NTA resin (Millipore) pre-equilibrated with lysis-buffer for 1 hour on ice with gentle shaking. The resin was washed with 3 separate 30 mM imidazole washes (15 mL) before elution with 250 mM imidazole. The elution fraction was dialyzed overnight at 4 °C into 7.3 mM HEPES pH 7.3, 200 mM NaCl and concentrated using an Amicon concentrator (Millipore-Sigma). Protein was then aliquoted, flash frozen in liquid nitrogen, and stored at - 80°C. FimX was expressed as described above and purified as previously described (30), except for the final purification step involving separation by size-exclusion chromatography (Superdex 200, Cytiva) equilibrated using 50 mM Tris-HCl pH 8.0, 250 mM NaCl, and 5 mM β-ME. Protein-containing fractions were determined using SDS-PAGE and then pooled for concentration using an Amicon concentrator.

### ATPase activity assay

All ATPase assays contained equimolar amount of ΔN1D-PilB or mutants (100 nM) to ensure valid comparison. PilB reaction buffer was prepared containing 50 mM HEPES pH 7.3, 200 mM NaCl, 5 mM MgCl_2_, 1 µM ZnCl_2_, and 1 mM ATP. Where applicable, the reaction buffer was supplemented with 1 µM cdGMP, and/or 200 nM of purified FimX. Aliquots of ΔN1D-PilB was thawed and the tube was centrifuged at maximum speed for 10 min at 4 °C to pellet insoluble precipitate. The remaining supernatant was transferred to a fresh tube and kept on ice. ATPase reactions were performed in a clear flat-bottom 96 well plate (Corning) in a total volume of 100 µL at 37 °C in a standing incubator. Following incubation and at the appropriate time point, 50 µL of reaction was mixed with 100 µL of BioMol green reagent (Enzo Lifesciences) incubated at room temperature for 15 min at which point absorbance at 620 nm was read with a spectrophotometer. Each ΔN1D-PilB reaction series was normalized to the timepoint at 0 min to ensure consistent background signal reduction. For overnight incubations, the 96-well plate lid was sealed with masking tape, and the reaction was performed as described with samples being normalized to an equivalent reaction well lacking PilB.

### Fluorescence microscopy experiments

T4P labeling experiments were performed using a cysteine knock-in method described previously, with some species-specific changes (4, 69). 100 µl of overnight cultures of *P. aeruginosa* were spotted onto prewarmed LB agar plates and grown upright at 37 °C for 4-5 hours. Plate-grown cells were then scraped off of plates using a P200 pipette tip and gently resuspended into 100 µl of fresh LB. AlexaFluor488-maleimide (Sigma) was added to the cell suspension for a final concentration of 50 ng/µl and incubated for 45-60 min in the dark at room temperature. Cells were washed once with 200 µl fresh LB using centrifugation settings of 18,000 x *g* for 1 min. Cell were resuspended into 200 µl of fresh LB. mNeonGreen-FimX strains were prepared exactly the same way except with gentamycin supplemented in the medium and with no AlexaFluor488-maleimide labeling steps. All imaging was performed under 1% agarose pads made with PBS solution. Cell bodies were imaged using phase-contrast microscopy on a Nikon Ti2-E microscope using a Plan Apo 100X oil immersion objective, a GFP/FITC/Cy2 filter set for pili and mNeonGreen-FimX, a Hamamatsu ORCA-Fusion Gen-III cCMOS camera, and Nikon NIS Elements Imaging Software. Measurements of pili were quantified manually using Fiji. Notably, populations of *P. aeruginosa* containing labeled T4P exhibit variability in labeling efficiency for unknown reasons. Therefore, T4P fluorescence signal contrast was adjusted so that T4P were clearly visible for each strain.

## Acknowledgements

We thank Dr. George O’Toole and Dr. Christopher Geiger for the cAMP reporter plasmid and their help in optimizing the detection assay. We thank Dr. Lynne Howell and Ian Yen for the *ydeH*-pBADGr plasmid. We also thank Dr. Sara Andres, Caitlin Doubleday, and the Andres lab for access to equipment and assistance in purifying all proteins used in this study. We thank the members of the Burrows lab for their helpful discussions and insights. This work was supported by a Canadian Institutes of Health Research Project grant (PJT-156080) to LLB and a National Institutes of Health grant (R35GM150916) to CKE. LLB holds a Tier 1 Canada Research Chair in Microbe-Surface Interactions (CRC 2021–00103). CKE is a Damon Runyon-Marilyn and Scott Urdang Breakthrough Scientist supported by the Damon Runyon Cancer Research Foundation (DFS6023). NR holds an NSERC CGS-D award. NY was funded through an NSERC USRA.

## SI Figure Captions

**Figure S1: Extension regulatory effector mutants produce no detectable surface pili.** Representative SDS-PAGE of sheared surface proteins. Major pilin subunits (PilA) are indicated. Sample loading was normalized to flagellin (FliC) levels.

**Figure S2: Mutations which suppress the *fimX* twitching deficit do not confer a growth benefit.** Bacterial growth curves across 18 hours in LB media. Points represent the means of triplicate samples from three independent experiments ± SD.

**Figure S3: Motility suppressor mutants remain susceptible to pilus-targeting phage PO4.** Bacterial growth curves across 18 hours in LB media with PO4 phage-challenge. Points represent the means of triplicate samples from three independent experiments ± SD.

**Figure S4: FimX complementation increases twitching motility in original Δ*fimX* suppressor isolates. (A)** Quantification of sub-agar stab twitching motility zones for Δ*fimX* twitching suppressor mutants complemented with FimX variants. Representative crystal violet-stained twitching zones are shown to the left. Scale bar = 1 cm. Bars represent the means of triplicate samples from three independent experiments ± SD. **(B)** Representative colonies showing pairwise interactions (pink) between FimX and FimX AAA mutant. PilZ was used as a known non-interacting negative control with FimX (52). Untagged T18/T25 plasmid and PilS-T18/PilS-T25 homodimers (63) were used as negative and positive controls respectively. **EV**: empty pHERD30T vector, ***fimX***: *fimX* in pHERD30T vector, **FimX AAA**: FimX AAA in pHERD30T vector.

**Figure S5: Expression of CyaB *in trans* bypasses the Pil-Chp network to increase twitching motility. (A)** Quantification of sub-agar stab twitching motility zones for a Δ*cyaB* mutant complemented with CyaB or a catalytically inactive mutant (D234S). Representative CyaB or CyaB D234S-expressing BTH101 colonies are shown in the inset to the right. Pink colour results from increased cAMP levels. Representative crystal violet-stained twitching zones are shown to the left. Bars represent the means of triplicate samples from three independent experiments ± SD. **(B)** Quantification of sub-agar stab twitching motility zones for a Δ*pilGH* double mutant complemented with CyaB. Representative twitching zones are shown to the left. Bars represent the means of triplicate samples from three independent experiments ± SD. All scale bars = 1 cm. **\*\*\***: *p* ≤ 0.001 (Two-tailed parametric *t*-test). **EV**: empty pHERD30T vector, **His_6_-CyaB (D234S)**: N-terminally hexa-histidine tagged CyaB or CyaB D234S in pHERD30T vector.

**Figure S6: Mutants with an mRuby3 cassette insertion upstream of the *pilMNOPQ* operon remain sensitive to pilus-targeting phage PO4.** Bacterial growth curves across 10 hours in LB media with PO4 phage-challenge. Points represent the means of triplicate samples from two independent experiments ± SD. **CyaB R456L**: CyaB R456L in pBADGr vector.

**Figure S7: Type II secretion system-dependent clearing on skim milk agar plates for original *fimX* suppressor isolates. (A)** Representative colonies and surrounding zones of clearance for the Δ*fimX* twitching suppressor mutants. Scale bar = 1 cm. **(B)** Quantification of the colony area subtracted from the skim milk clearance area. Bars represent the means of duplicate samples from three independent experiments ± SD. **\***: 0.05 ≥ *p* ≥ 0.01; **\*\*\***: 0.001 ≥ *p* (Two-tailed parametric *t*-test).

**Figure S8: Alphafold3-predicted models of PilB supressor mutants *pilB* Δ1bp and PilB T430P. (A)** Model of *P. aeruginosa* PilB (PilB^Pa^) monomer. The N1D and linker region (dark grey), N2D (light blue), and CTD (white) are indicated. The *pilB* Δ1bp C-terminal extension is shown in dark blue. PilB T430 side chain is shown in stick and in orange. The model PAE plot is shown to the right. **(B)** *pilB* Δ1bp full length homohexameric Alphafold3 predicted model is shown below with the extended sequence residues highligted in dark blue. The predicted PilC interface is shown on the bottom. The N1Ds and linker regions for each monomer are hidden. The model on the bottom has the two front monomers hidden for clarity. **(C)** X-ray crystal structure of hexameric PilB from *G. metallireducens* (5TSH) with sequence aligned residues of interest N429 (PilB^Pa^ T430) and R430 (PilB^Pa^ R431) highlighted in orange and magenta, respectively. C2-symetric monmers are highlited in the same colours. Dashed lines indicate approximate protomer interfaces.

**Figure S9: *pilB* Δ1bp constructs complement twitching motility to at least WT levels. (A)** Quantification of sub-agar stab twitching zone areas of a *pilB*::Tn5 mutant overexpressing *pilB* Δ1bp 3ʹ truncations. Representative crystal violet-stained twitching zones are shown to the left. PilB expression was induced with 0.05% arabinose. Bars represent the means of duplicate samples from three independent experiments ± SD. **(B)** Quantification of sub-agar stab twitching zone areas of WT cells overexpressing *pilB* Δ1bp 3ʹ truncations. Representative twitching zones are shown to the left. PilB expression was induced with 0.05% arabinose. Bars represent the means of duplicate samples from three independent experiments ± SD. **EV**: empty pHERD30T vector, ***pilB* Δ1bp-3 to 21**: *pilB* Δ1bp-3 to 21 in pHERD30T vector. All scale bars = 1 cm. **ns**: *p* ≥ 0.05; **\*\***: 0.01 ≥ *p* ≥ 0.001 (Two-tailed parametric *t*-test).

**Figure S10: PilB T430X constructs complement twitching motility to at least WT levels. (A)** Quantification of sub-agar stab twitching zone areas of a *pilB*::Tn5 mutant overexpressing T430X mutants. Representative crystal violet-stained twitching zones are shown to the left. PilB expression was induced with 0.05% arabinose. Bars represent the means of duplicate samples from three independent experiments ± SD. **(B)** Quantification of sub-agar stab twitching zone areas of WT cells overexpressing T430X mutants. Representative twitching zones are shown to the left. PilB expression was induced with 0.05% arabinose. Bars represent the means of duplicate samples from three independent experiments ± SD. **PilB T430X**: indicated PilB substitution in pHERD30T vector. All scale bars = 1 cm. **ns**: *p* ≥ 0.05; **\***: 0.05 ≥ *p* ≥ 0.01 (Two-tailed parametric *t*-test).

**Figure S11: PilB T429/431X constructs complement twitching motility. (A)** Quantification of sub-agar stab twitching zone areas of a *pilB*::Tn5 mutant overexpressing T429/431X mutants. Representative crystal violet-stained twitching zones are shown to the left. PilB expression was induced with 0.05% arabinose. Bars represent the means of duplicate samples from three independent experiments ± SD. **(B)** Quantification of sub-agar stab twitching zone areas of WT cells overexpressing T429/431X mutants. Representative twitching zones are shown to the left. PilB expression was induced with 0.05% arabinose. Bars represent the means of duplicate samples from three independent experiments ± SD. **PilB T429/431X**: indicated PilB substitution in pHERD30T vector. All scale bars = 1 cm. **ns**: *p* ≥ 0.05; **\*\*\***: 0.001 ≥ *p* (Two-tailed parametric *t*-test).

**Figure S12: Purified ΔN1D-His_6_-PilB and FimX protein samples.** SDS-PAGE of representative protein samples used in the study. Samples are diluted to the same concentrations added to the reaction mixture preparation.

**Figure S13: PilA A86C mutants retain twitching motility.** Quantification of sub-agar stab twitching zone areas of PilA A86C mutants. Representative crystal violet stained twitching zones are shown to the left. Bars represent the means of triplicate samples from three independent experiments ± SD. Scale bar = 1 cm.

**Figure S14: Vector control assays for FimX localization studies. (A)** Quantification of sub-agar stab twitching zone areas of Δ*fimX* mutants complemented with N-terminal fusion of mNeonGreen (mNGr) to FimX. Representative crystal violet stained twitching zones are shown to the left. Expression was not induced with arabinose. Bars represent the means of triplicate samples from three independent experiments ± SD. Scale bar = 1 cm. **EV**: empty pHERD30T, **mNGr-FimX (AAA)**: N-terminal fusion of mNeonGreen to FimX or FimX AAA in pHERD30T vector. **(B)** Representative images of Δ*fimX* mutant cells carrying empty pHERD30T with background fluorescence subtracted. Scale bar = 2 µm. **\*\*\***: 0.001 ≥ *p* (Two-tailed parametric *t*-test).

**Figure S15: PilB mutations contributing to enhanced twitching motility in Δ*fimX* are rare. (A)** Total protein sequence length of PilB orthologues in *P. aeruginosa* from the *Pseudomonas*.com database. Sequences which are the same length as PAO1 PilB (566 residues) are indicated in blue. **(B)** Percentage of sequences with an identifiable α-helix-disrupting residue ± three amino acids from the position which aligns with PAO1 PilB T430.

