## supplementary data for "Twitching motility suppressors reveal a role for FimX in type IV pilus extension dynamics"

**Supplementary Figures S1-S15 and Captions for Roberge et al., Twitching motility suppressors reveal a role for FimX in type IV pilus extension dynamics.**

### SI Figure Captions

**Figure S1: Extension regulatory effector mutants produce no detectable surface pili.** Representative SDS-PAGE of sheared surface proteins. Major pilin subunits (PilA) are indicated. Sample loading was normalized to flagellin (FliC) levels.

**Figure S2: Mutations which suppress the *fimX* twitching deficit do not confer a growth benefit.** Bacterial growth curves across 18 hours in LB media. Points represent the means of triplicate samples from three independent experiments ± SD.

**Figure S3: Motility suppressor mutants remain susceptible to pilus-targeting phage PO4.** Bacterial growth curves across 18 hours in LB media with PO4 phage-challenge. Points represent the means of triplicate samples from three independent experiments ± SD.

**Figure S4: FimX complementation increases twitching motility in original Δ*fimX* suppressor isolates. (A)** Quantification of sub-agar stab twitching motility zones for Δ*fimX* twitching suppressor mutants complemented with FimX variants. Representative crystal violet-stained twitching zones are shown to the left. Scale bar = 1 cm. Bars represent the means of triplicate samples from three independent experiments ± SD. **(B)** Representative colonies showing pairwise interactions (pink) between FimX and FimX AAA mutant. PilZ was used as a known non-interacting negative control with FimX (1). Untagged T18/T25 plasmid and PilS-T18/PilS-T25 homodimers (2) were used as negative and positive controls respectively. **EV**: empty pHERD30T vector, ***fimX***: *fimX* in pHERD30T vector, **FimX AAA**: FimX AAA in pHERD30T vector.

**Figure S5: Expression of CyaB *in trans* bypasses the Pil-Chp network to increase twitching motility. (A)** Quantification of sub-agar stab twitching motility zones for a Δ*cyaB* mutant complemented with CyaB or a catalytically inactive mutant (D234S). Representative CyaB or CyaB D234S-expressing BTH101 colonies are shown in the inset to the right. Pink colour results from increased cAMP levels. Representative crystal violet-stained twitching zones are shown to the left. Bars represent the means of triplicate samples from three independent experiments ± SD. **(B)** Quantification of sub-agar stab twitching motility zones for a Δ*pilGH* double mutant complemented with CyaB. Representative twitching zones are shown to the left. Bars represent the means of triplicate samples from three independent experiments ± SD. All scale bars = 1 cm. ***: $p \leq 0.001$ (Two-tailed parametric *t*-test). **EV**: empty pHERD30T vector, **His$_6$-CyaB (D234S)**: N-terminally hexa-histidine tagged CyaB or CyaB D234S in pHERD30T vector.

**Figure S6: Mutants with an mRuby3 cassette insertion upstream of the *pilMNOPQ* operon remain sensitive to pilus-targeting phage PO4.** Bacterial growth curves across 10 hours in LB media with PO4 phage-challenge. Points represent the means of triplicate samples from two independent experiments ± SD. **CyaB R456L**: CyaB R456L in pBADGr vector.

**Figure S7: Type II secretion system-dependent clearing on skim milk agar plates for original *fimX* suppressor isolates. (A)** Representative colonies and surrounding zones of clearance for the Δ*fimX* twitching suppressor mutants. Scale bar = 1 cm. **(B)** Quantification of the colony area subtracted from the skim milk clearance area. Bars represent the means of duplicate samples from three independent experiments ± SD. *: $0.05 \geq p \geq 0.01$; ***: $0.001 \geq p$ (Two-tailed parametric *t*-test).

**Figure S8: Alphafold3-predicted models of PilB supressor mutants *pilB* Δ1bp and PilB T430P. (A)** Model of *P. aeruginosa* PilB (PilB$^{Pa}$) monomer. The N1D and linker region (dark grey), N2D (light blue), and CTD (white) are indicated. The *pilB* Δ1bp C-terminal extension is shown in dark blue. PilB T430 side chain is shown in stick and in orange. The model PAE plot is shown to the right. **(B)** *pilB* Δ1bp full length homohexameric Alphafold3 predicted model is shown below with the extended sequence residues highligted in dark blue. The predicted PilC interface is shown on the bottom. The N1Ds and linker regions for each monomer are hidden. The model on the bottom has the two front monomers hidden for clarity. **(C)** X-ray crystal structure of hexameric PilB from *G. metallireducens* (5TSH) with sequence aligned residues of interest N429 (PilB$^{Pa}$ T430) and R430 (PilB$^{Pa}$ R431) highlighted in orange and magenta, respectively. C2-symetric monmers are highlited in the same colours. Dashed lines indicate approximate protomer interfaces.

**Figure S9: *pilB* Δ1bp constructs complement twitching motility to at least WT levels. (A)** Quantification of sub-agar stab twitching zone areas of a *pilB*::Tn5 mutant overexpressing *pilB* Δ1bp 3′ truncations. Representative crystal violet-stained twitching zones are shown to the left. PilB expression was induced with 0.05% arabinose. Bars represent the means of duplicate samples from three independent experiments ± SD. **(B)** Quantification of sub-agar stab twitching zone areas of WT cells overexpressing *pilB* Δ1bp 3′ truncations. Representative twitching zones are shown to the left. PilB expression was induced with 0.05% arabinose. Bars represent the means of duplicate samples from three independent experiments ± SD. **EV**: empty pHERD30T vector, ***pilB* Δ1bp-3 to 21**: *pilB* Δ1bp-3 to 21 in pHERD30T vector. All scale bars = 1 cm. **ns**: $p \geq 0.05$; **: $0.01 \geq p \geq 0.001$ (Two-tailed parametric *t*-test).

**Figure S10: PilB T430X constructs complement twitching motility to at least WT levels. (A)** Quantification of sub-agar stab twitching zone areas of a *pilB*::Tn5 mutant overexpressing T430X mutants. Representative crystal violet-stained twitching zones are shown to the left. PilB expression was induced with 0.05% arabinose. Bars represent the means of duplicate samples from three independent experiments ± SD. **(B)** Quantification of sub-agar stab twitching zone areas of WT cells overexpressing T430X mutants. Representative twitching zones are shown to the left. PilB expression was induced with 0.05% arabinose. Bars represent the means of duplicate samples from three independent experiments ± SD. **PilB T430X**: indicated PilB substitution in pHERD30T vector. All scale bars = 1 cm. **ns**: $p \geq 0.05$; *: $0.05 \geq p \geq 0.01$ (Two-tailed parametric *t*-test).

**Figure S11: PilB T429/431X constructs complement twitching motility. (A)** Quantification of sub-agar stab twitching zone areas of a *pilB*::Tn5 mutant overexpressing T429/431X mutants. Representative crystal violet-stained twitching zones are shown to the left. PilB expression was induced with 0.05% arabinose. Bars represent the means of duplicate samples from three independent experiments ± SD. **(B)** Quantification of sub-agar stab twitching zone areas of WT cells overexpressing T429/431X mutants. Representative twitching zones are shown to the left. PilB expression was induced with 0.05% arabinose. Bars represent the means of duplicate samples from three independent experiments ± SD. **PilB T429/431X**: indicated PilB substitution in pHERD30T vector. All scale bars = 1 cm. **ns**: $p \geq 0.05$; ***: $0.001 \geq p$ (Two-tailed parametric *t*-test).

**Figure S12: Purified ΔN1D-His$_6$-PilB and FimX protein samples.** SDS-PAGE of representative protein samples used in the study. Samples are diluted to the same concentrations added to the reaction mixture preparation.

**Figure S13: PilA A86C mutants retain twitching motility.** Quantification of sub-agar stab twitching zone areas of PilA A86C mutants. Representative crystal violet stained twitching zones are shown to the left. Bars represent the means of triplicate samples from three independent experiments ± SD. Scale bar = 1 cm.

**Figure S14: Vector control assays for FimX localization studies. (A)** Quantification of sub-agar stab twitching zone areas of Δ*fimX* mutants complemented with N-terminal fusion of mNeonGreen (mNGr) to FimX. Representative crystal violet stained twitching zones are shown to the left. Expression was not induced with arabinose. Bars represent the means of triplicate samples from three independent experiments ± SD. Scale bar = 1 cm. **EV**: empty pHERD30T, **mNGr-FimX (AAA)**: N-terminal fusion of mNeonGreen to FimX or FimX AAA in pHERD30T vector. **(B)** Representative images of Δ*fimX* mutant cells carrying empty pHERD30T with background fluorescence subtracted. Scale bar = 2 µm. ***: $0.001 \geq p$ (Two-tailed parametric *t*-test).

**Figure S15: PilB mutations contributing to enhanced twitching motility in Δ*fimX* are rare.** **(A)** Total protein sequence length of PilB orthologues in *P. aeruginosa* from the *Pseudomonas*.com database. Sequences which are the same length as PAO1 PilB (566 residues) are indicated in blue. **(B)** Percentage of sequences with an identifiable α-helix-disrupting residue ± three amino acids from the position which aligns with PAO1 PilB T430.

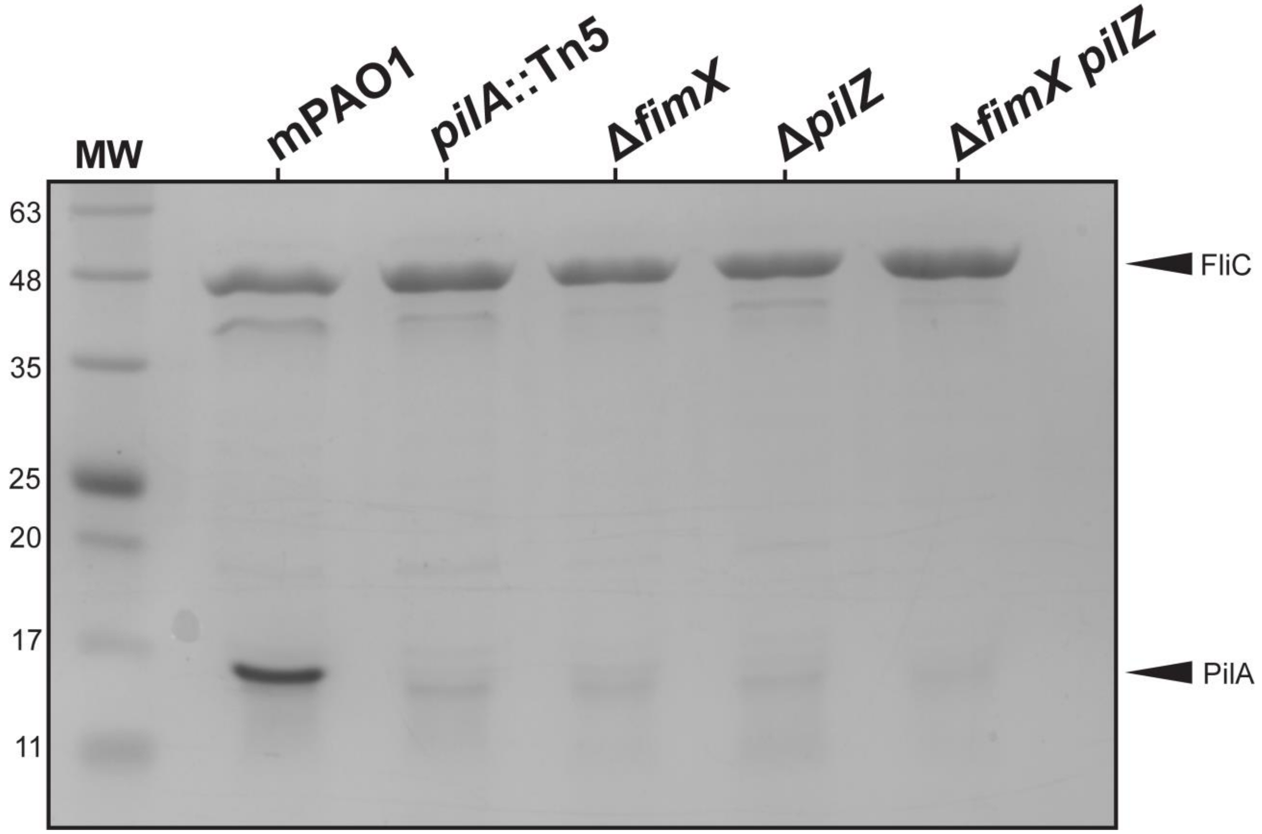
MW
mPAO1
pilA::Tn5
ΔfimX
ΔpilZ
ΔfimX pilZ
63
48
35
25
20
17
11
FliC
PilA

Fig S1

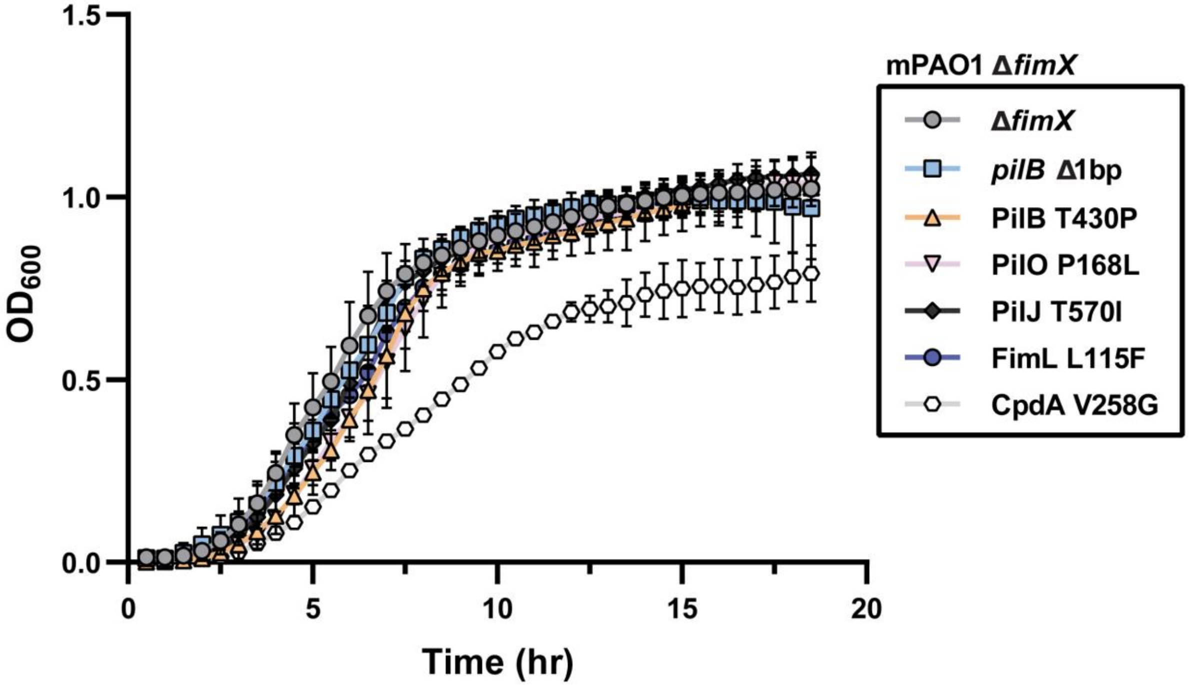
1.5
1.0
0.5
0.0
$OD_{600}$
0
5
10
15
20
Time (hr)
mPAO1 ΔfimX
ΔfimX
pilB Δ1bp
PilB T430P
PilO P168L
PilJ T570I
FimL L115F
CpdA V258G

Fig S2

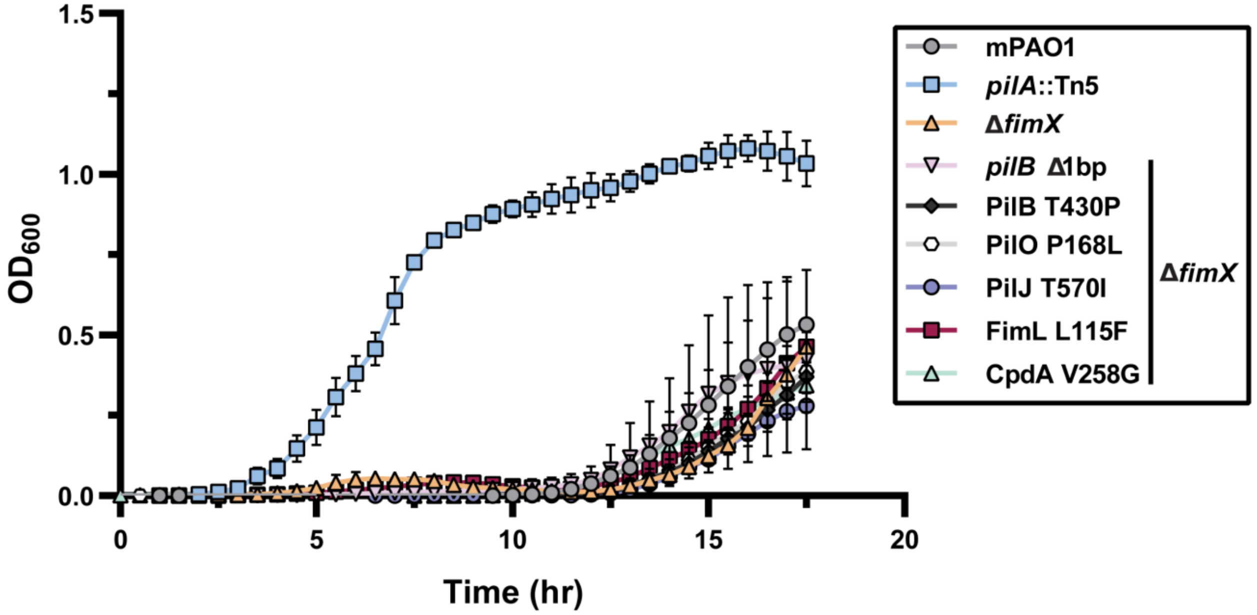
1.5
1.0
0.5
0.0
OD$_{600}$
0
5
10
15
20
Time (hr)
mPAO1
*pilA*::Tn5
Δ*fimX*
*pilB* Δ1bp
PilB T430P
PilO P168L
PilJ T570I
FimL L115F
CpdA V258G
Δ*fimX*

**Fig S3**

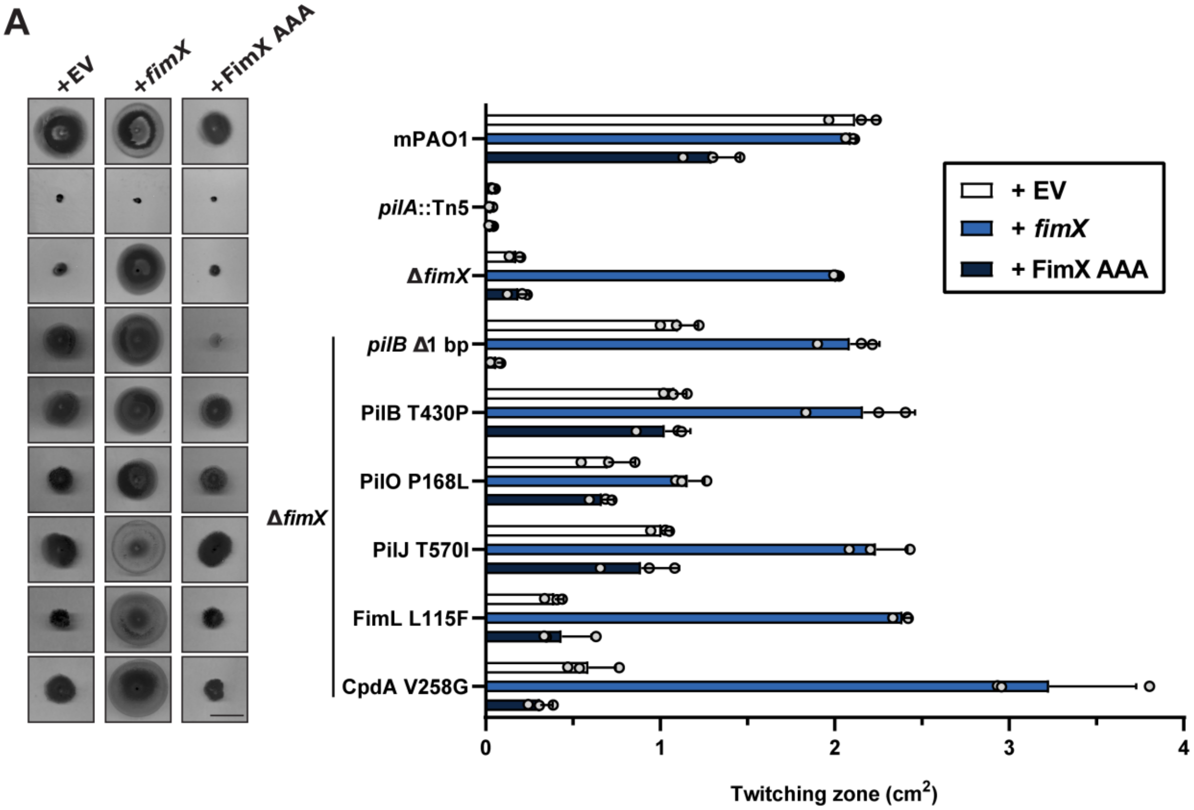
A
+EV
+fimX
+FimX AAA
mPAO1
pilA::Tn5
ΔfimX
pilB Δ1 bp
PilB T430P
PilO P168L
PilJ T570I
FimL L115F
CpdA V258G
ΔfimX
+ EV
+ fimX
+ FimX AAA
0
1
2
3
4
Twitching zone (cm2)

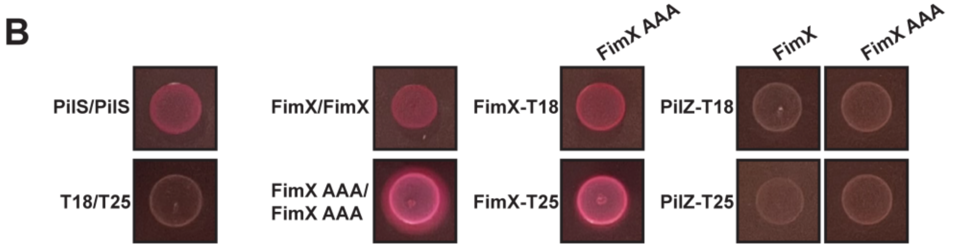
B
PilS/PilS
T18/T25
FimX/FimX
FimX AAA/
FimX AAA
FimX AAA
FimX-T18
FimX-T25
FimX
FimX AAA
PilZ-T18
PilZ-T25

Fig S4

A

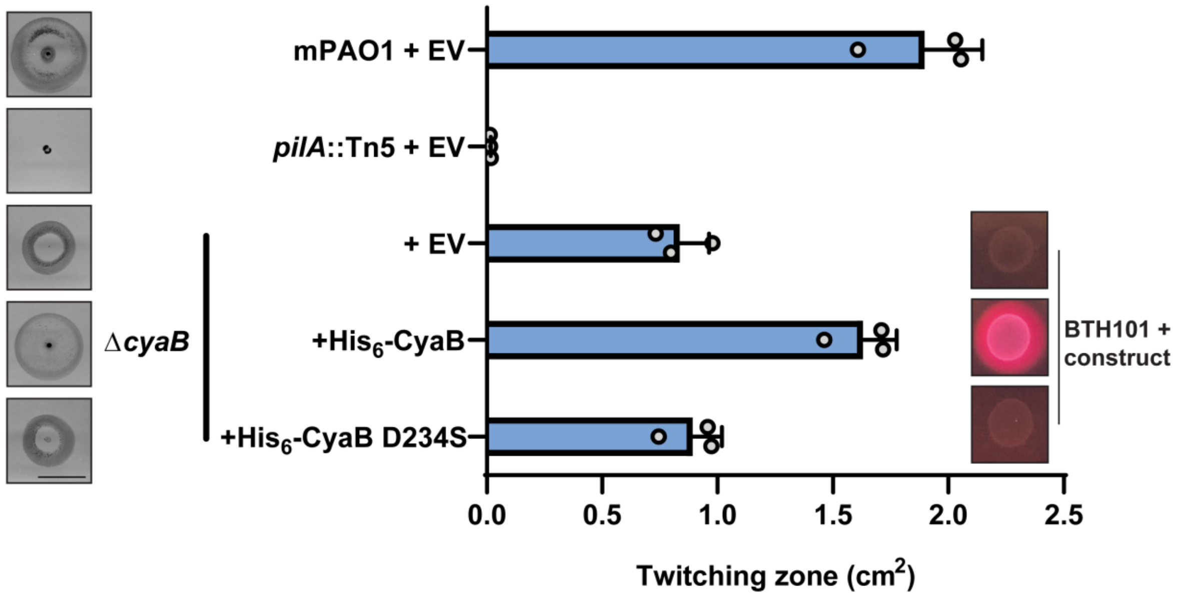
mPAO1 + EV
pilA::Tn5 + EV
ΔcyaB
+ EV
+His6-CyaB
+His6-CyaB D234S
BTH101 +
construct
0.0
0.5
1.0
1.5
2.0
2.5
Twitching zone (cm2)

B

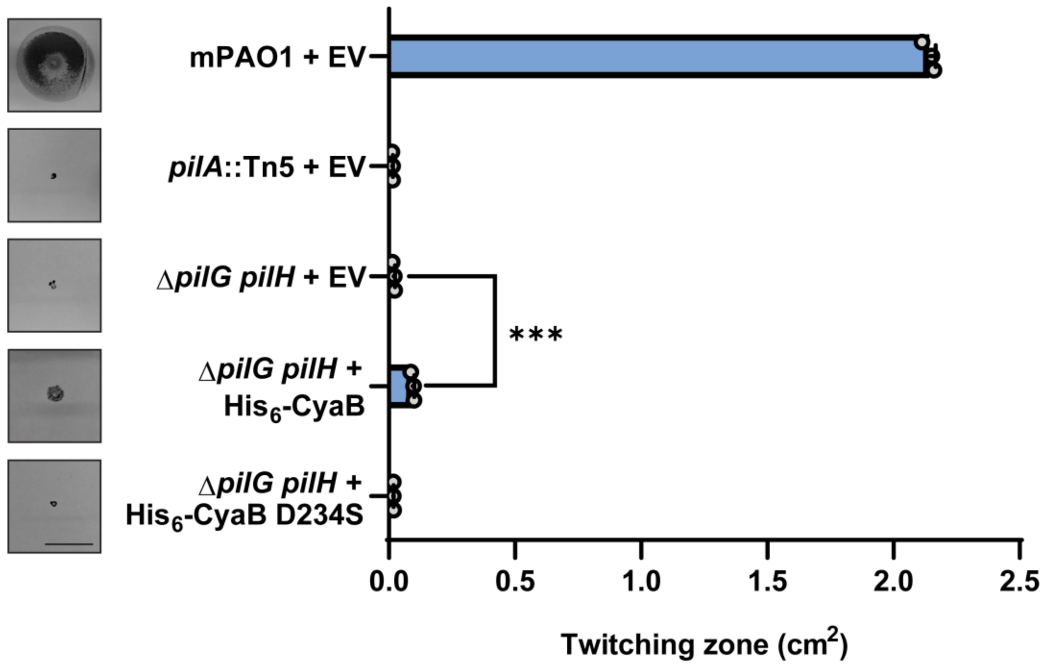
mPAO1 + EV
pilA::Tn5 + EV
ΔpilG pilH + EV
ΔpilG pilH +
His6-CyaB
ΔpilG pilH +
His6-CyaB D234S
***
0.0
0.5
1.0
1.5
2.0
2.5
Twitching zone (cm2)

Fig S5

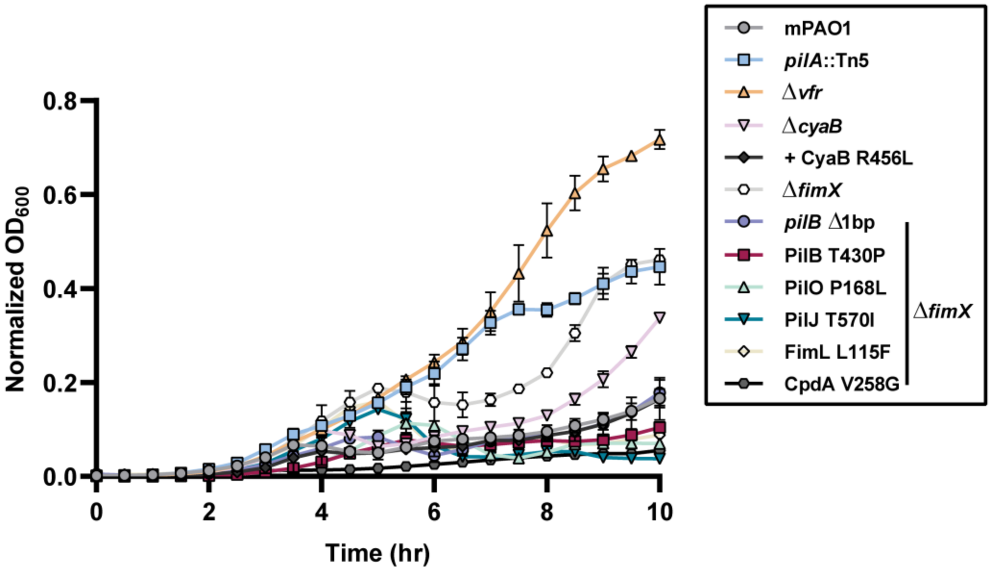
Normalized $OD_{600}$
0.8
0.6
0.4
0.2
0.0
0
2
4
6
8
10
Time (hr)
mPAO1
*pilA*::Tn5
Δ*vfr*
Δ*cyaB*
+ CyaB R456L
Δ*fimX*
*pilB* Δ1bp
PilB T430P
PilO P168L
PilJ T570I
FimL L115F
CpdA V258G
Δ*fimX*

**Fig S6**

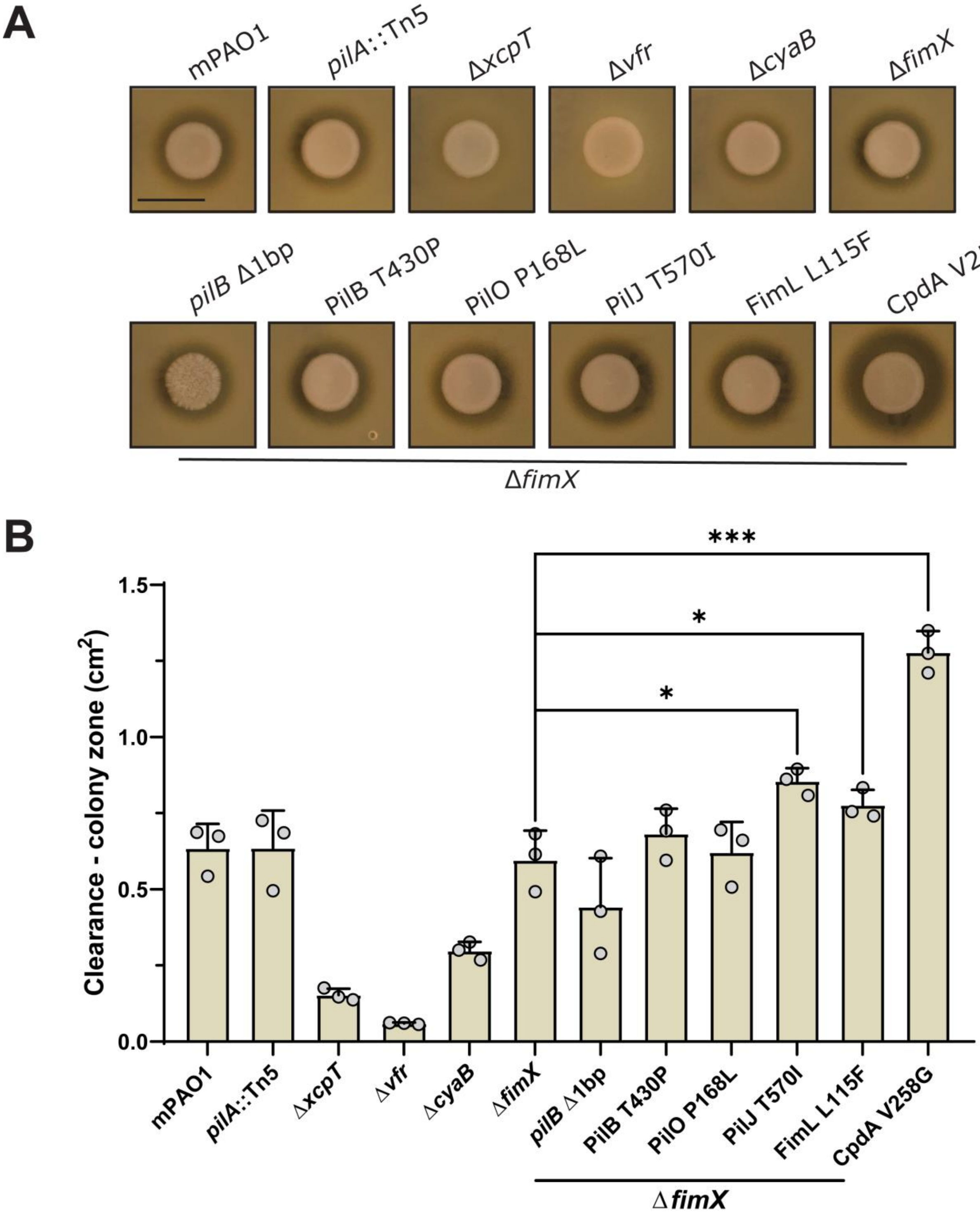
A
mPAO1
pilA::Tn5
ΔxcpT
Δvfr
ΔcyaB
ΔfimX
pilB Δ1bp
PilB T430P
PilO P168L
PilJ T570I
FimL L115F
CpdA V258G
ΔfimX
B
***
*
*
Clearance - colony zone (cm2)
1.5
1.0
0.5
0.0
mPAO1
pilA::Tn5
ΔxcpT
Δvfr
ΔcyaB
ΔfimX
pilB Δ1bp
PilB T430P
PilO P168L
PilJ T570I
FimL L115F
CpdA V258G
ΔfimX

**Fig S7**

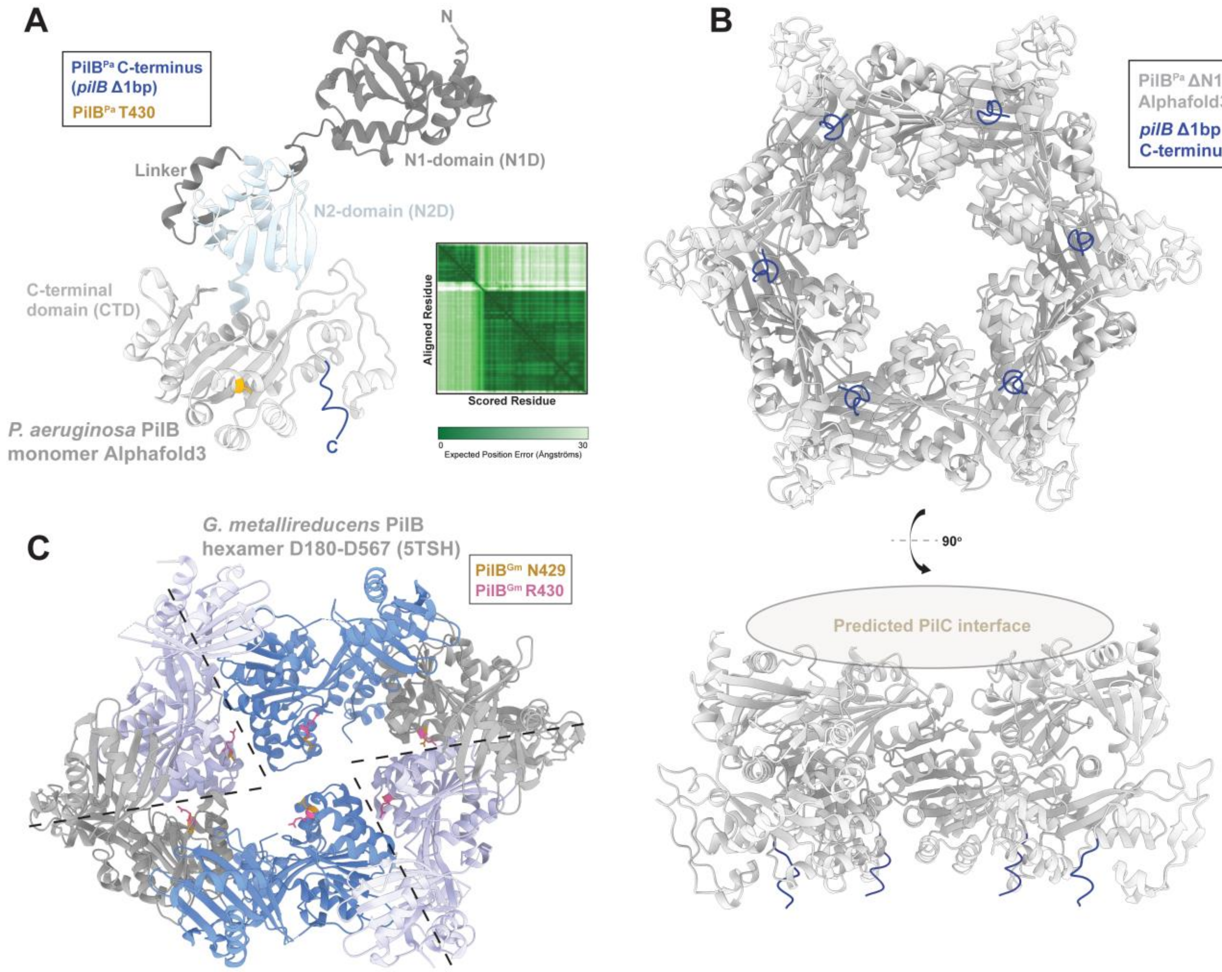
A
PilB^Pa C-terminus (pilB Δ1bp)
PilB^Pa T430
N
N1-domain (N1D)
Linker
N2-domain (N2D)
C-terminal domain (CTD)
Aligned Residue
Scored Residue
0
30
Expected Position Error (Ångströms)
P. aeruginosa PilB monomer Alphafold3
C
B
PilB^Pa ΔN1D Alphafold3
pilB Δ1bp C-terminus
90°
Predicted PilC interface
C
G. metallireducens PilB hexamer D180-D567 (5TSH)
PilB^Gm N429
PilB^Gm R430

**Fig S8**

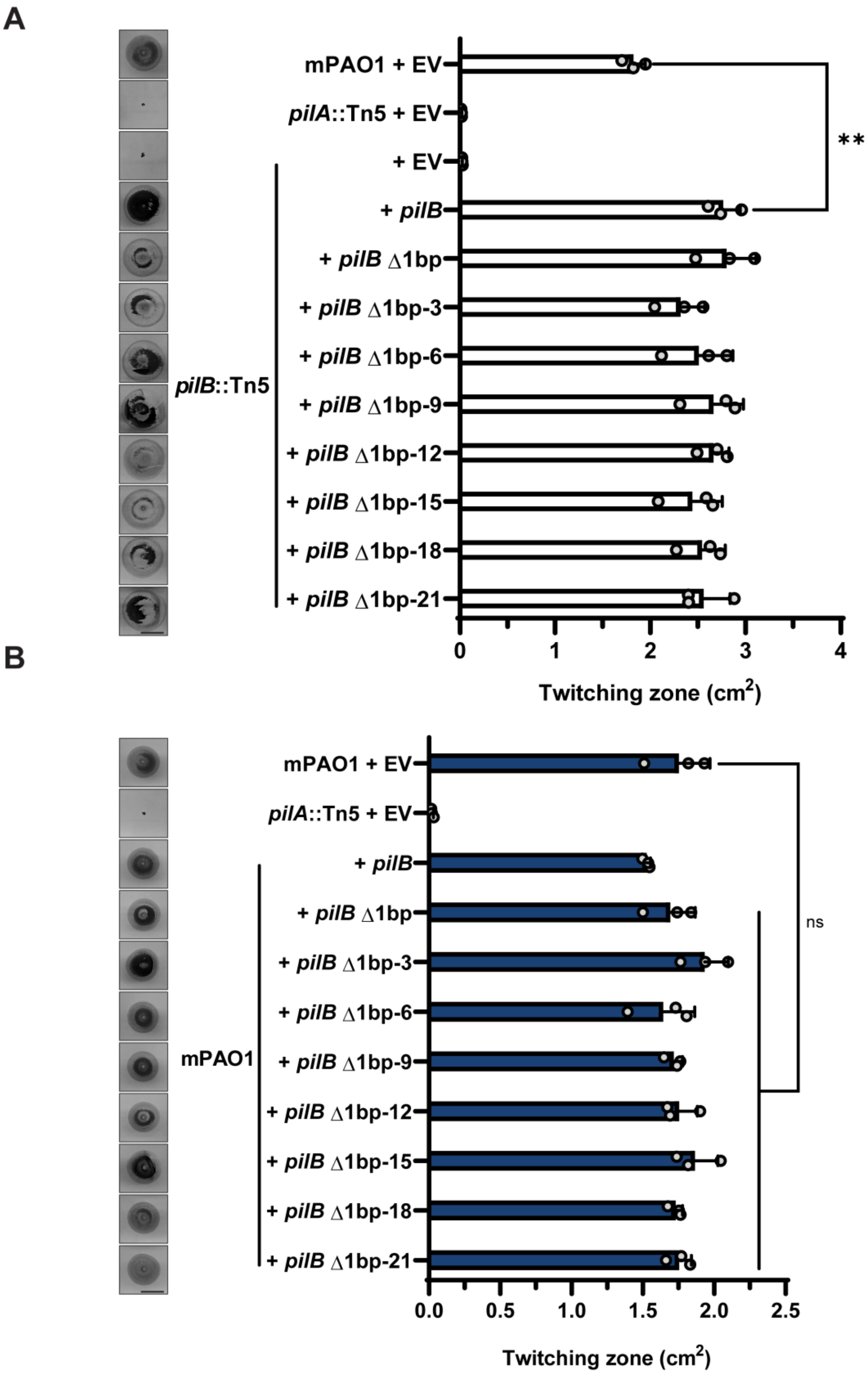
A
mPAO1 + EV
pilA::Tn5 + EV
+ EV
+ pilB
+ pilB Δ1bp
+ pilB Δ1bp-3
+ pilB Δ1bp-6
+ pilB Δ1bp-9
+ pilB Δ1bp-12
+ pilB Δ1bp-15
+ pilB Δ1bp-18
+ pilB Δ1bp-21
pilB::Tn5
**
0
1
2
3
4
Twitching zone (cm2)
B
mPAO1 + EV
pilA::Tn5 + EV
+ pilB
+ pilB Δ1bp
+ pilB Δ1bp-3
+ pilB Δ1bp-6
+ pilB Δ1bp-9
+ pilB Δ1bp-12
+ pilB Δ1bp-15
+ pilB Δ1bp-18
+ pilB Δ1bp-21
mPAO1
ns
0.0
0.5
1.0
1.5
2.0
2.5
Twitching zone (cm2)

**Fig S9**

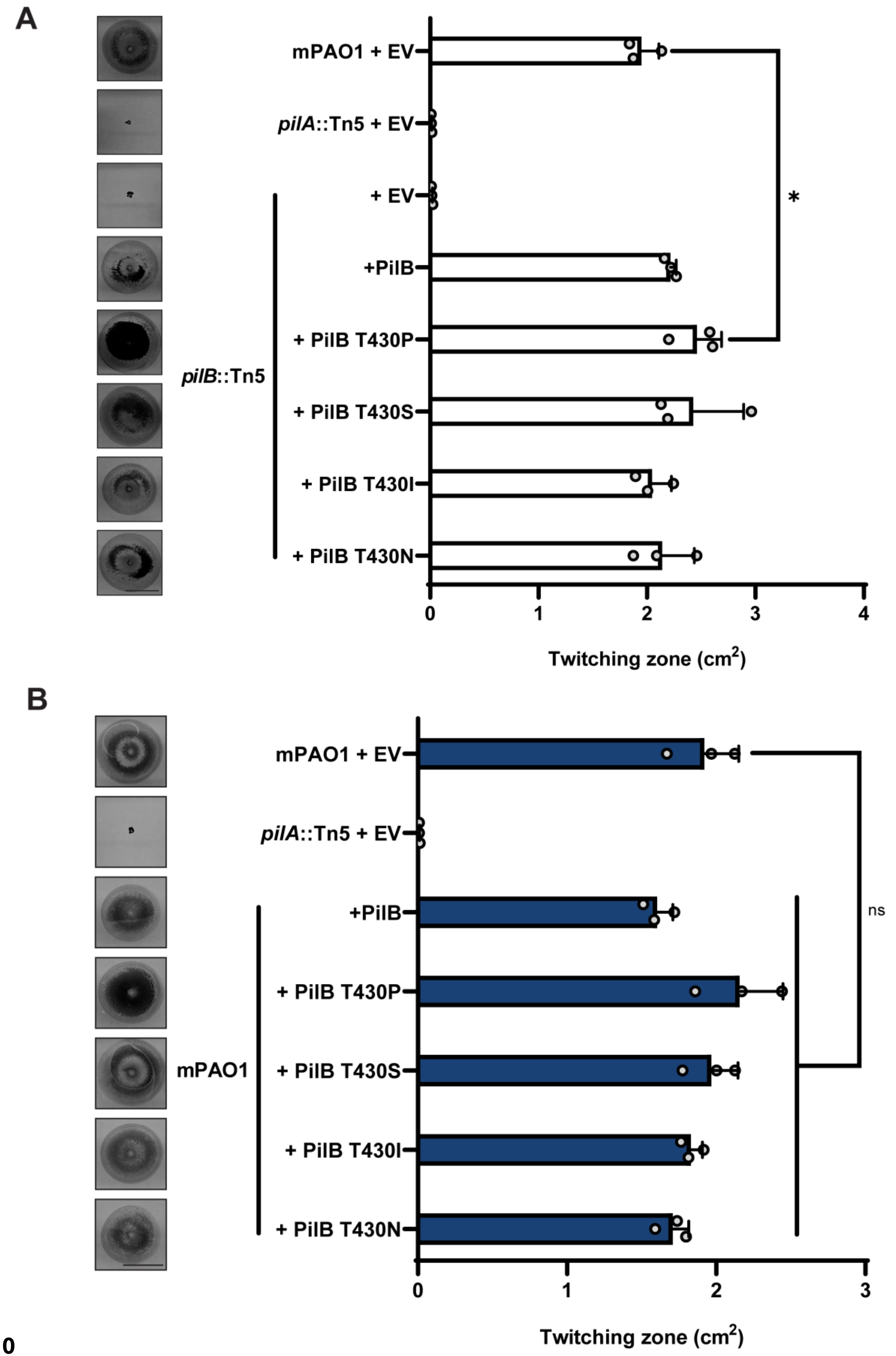
A
mPAO1 + EV
pilA::Tn5 + EV
+ EV
+PilB
+ PilB T430P
+ PilB T430S
+ PilB T430I
+ PilB T430N
pilB::Tn5
*
0
1
2
3
4
Twitching zone (cm2)
B
mPAO1 + EV
pilA::Tn5 + EV
+PilB
+ PilB T430P
+ PilB T430S
+ PilB T430I
+ PilB T430N
mPAO1
ns
0
1
2
3
Twitching zone (cm2)

Fig S10

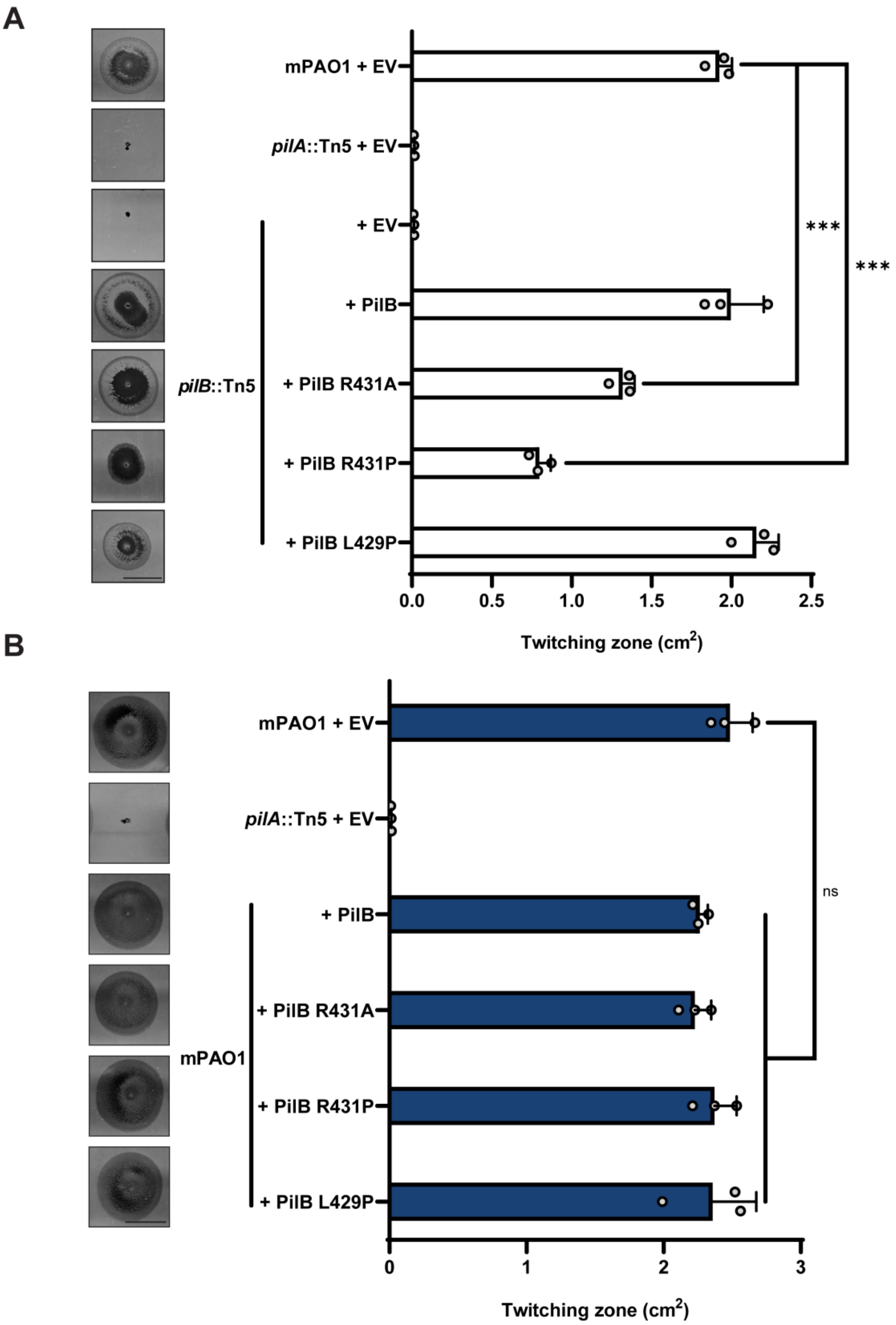

A
mPAO1 + EV
pilA::Tn5 + EV
+ EV
+ PilB
+ PilB R431A
+ PilB R431P
+ PilB L429P
pilB::Tn5
***
***
0.0
0.5
1.0
1.5
2.0
2.5
Twitching zone (cm2)
B
mPAO1 + EV
pilA::Tn5 + EV
+ PilB
+ PilB R431A
+ PilB R431P
+ PilB L429P
mPAO1
ns
0
1
2
3
Twitching zone (cm2)


**Fig S11**

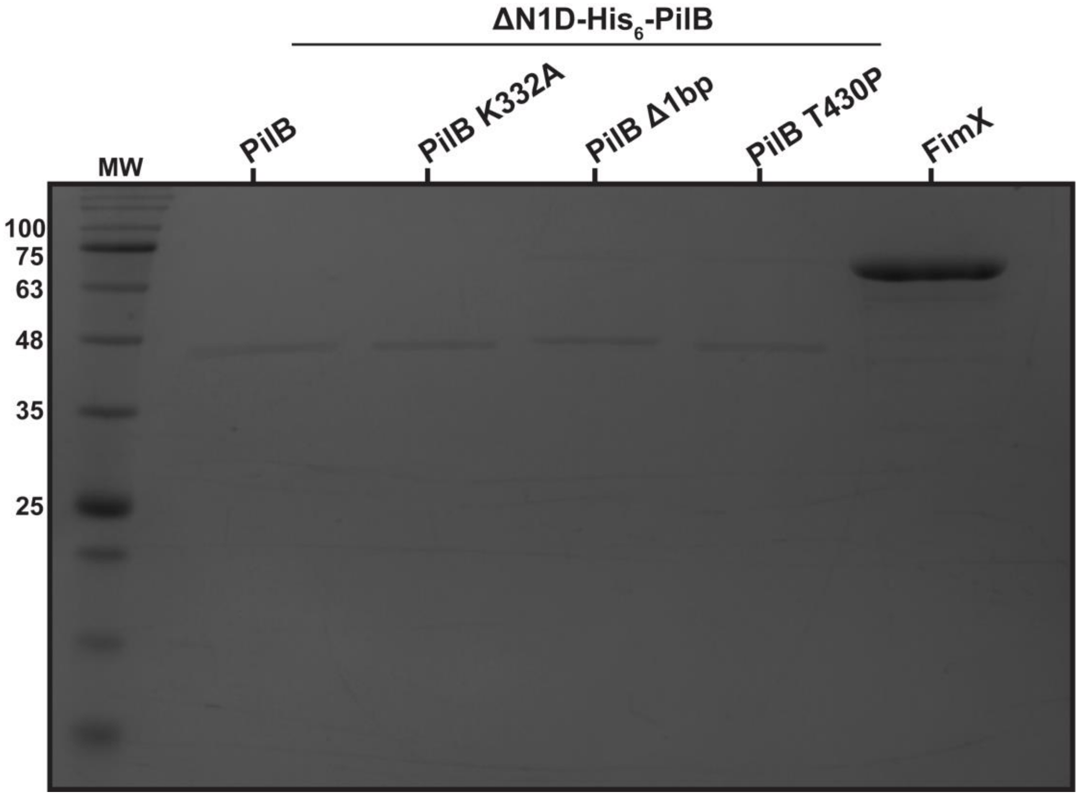


Fig S12

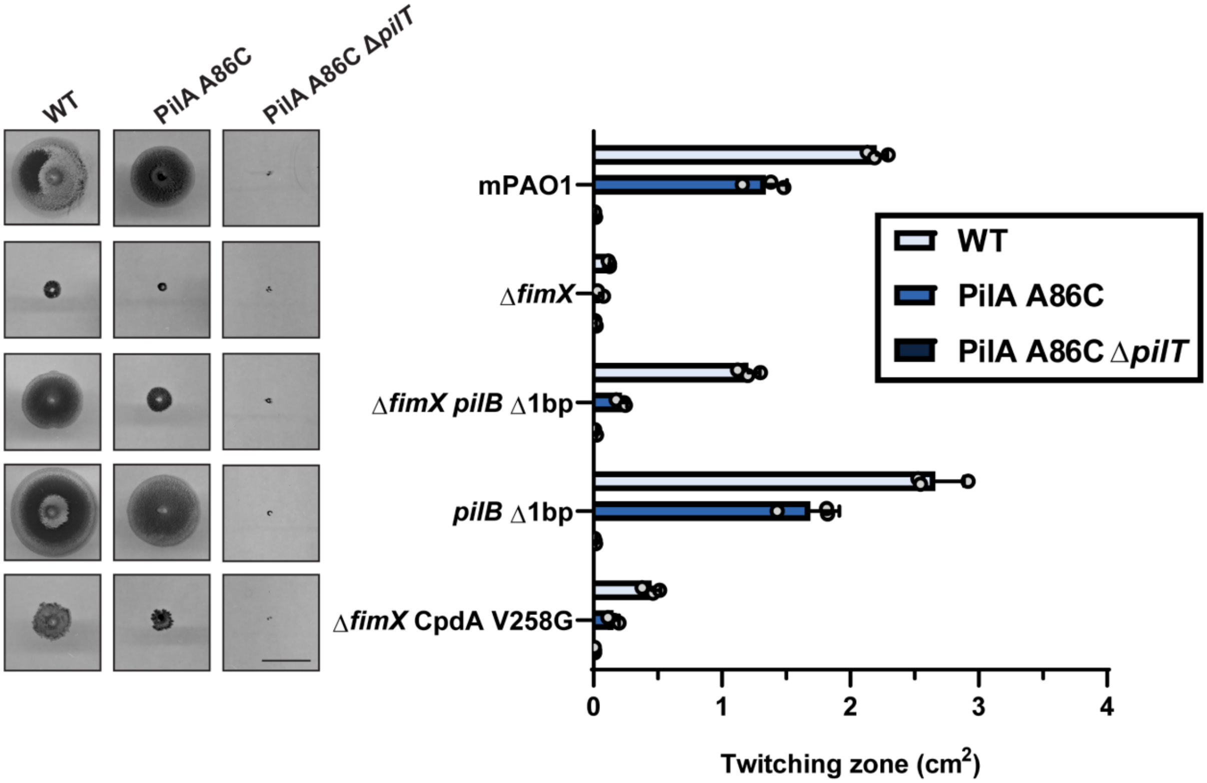


Fig S13

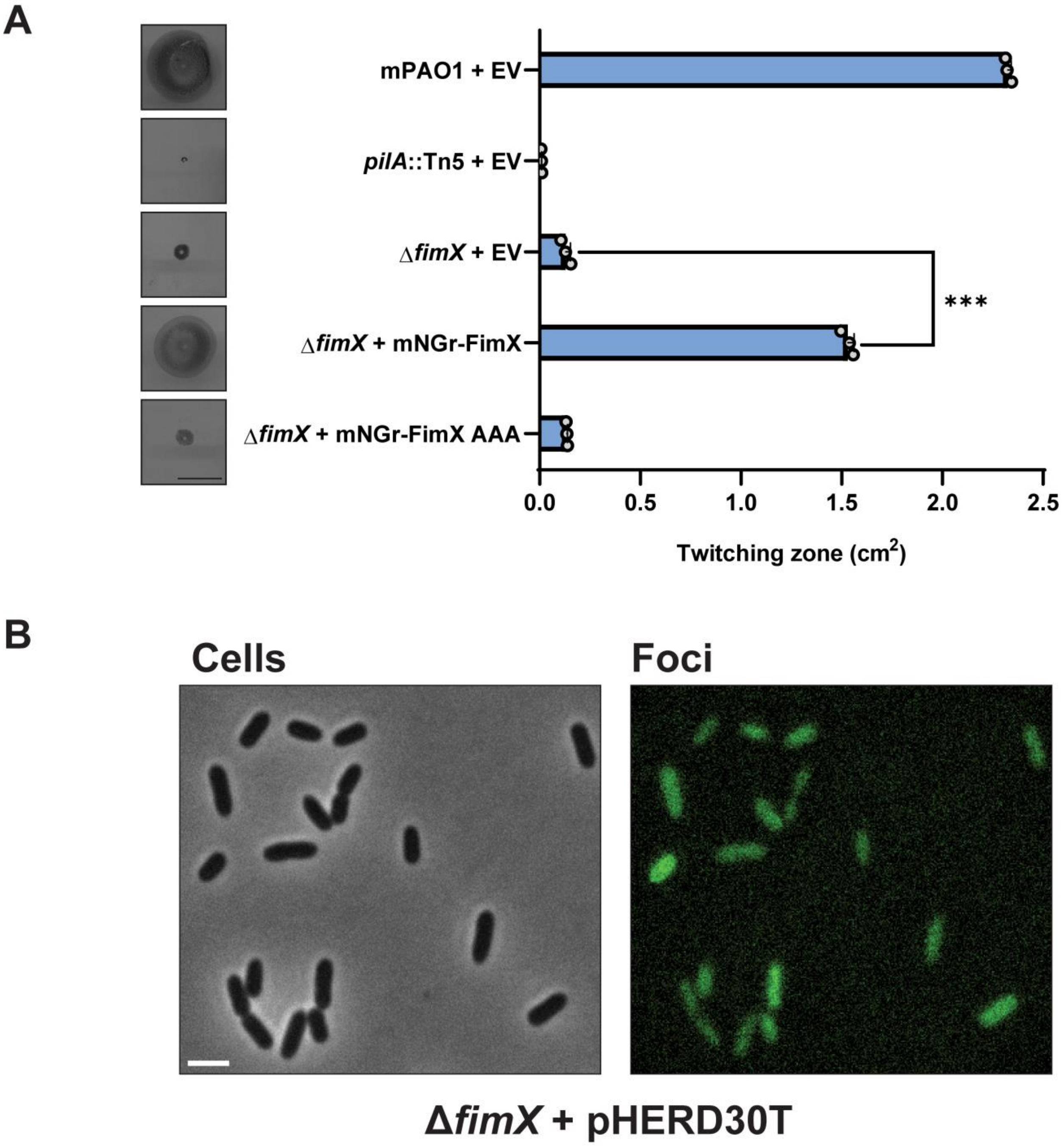
A
mPAO1 + EV
pilA::Tn5 + EV
ΔfimX + EV
ΔfimX + mNGr-FimX
ΔfimX + mNGr-FimX AAA
***
0.0
0.5
1.0
1.5
2.0
2.5
Twitching zone (cm2)
B
Cells
Foci
ΔfimX + pHERD30T

**Fig S14**

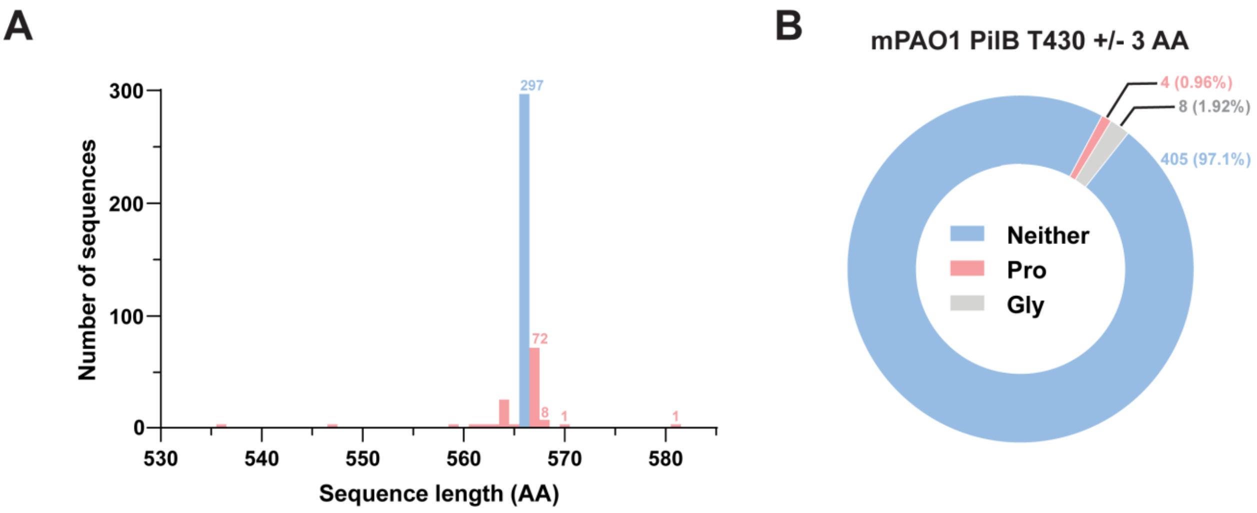
A
300
200
100
0
Number of sequences
297
72
8
1
1
530
540
550
560
570
580
Sequence length (AA)
B
mPAO1 PilB T430 +/- 3 AA
4 (0.96%)
8 (1.92%)
405 (97.1%)
Neither
Pro
Gly

**Fig S15**

**SI Tables for Roberge et al., Twitching motility suppressors reveal a role for FimX in type IV pilus extension dynamics.**

**Table S1: All putative suppressor mutations identified in this study.**

| Gene | Mutation | Function | Validated? |
|---|---|---|---|
| | **PilB** | | |
| *pilB* | Δ1bp – A1697 | Extension ATPase | Y |
| *pilB* | T430P | - | Y |
| *pilB* | D181N | - | N |
| *pilB* | E558D | - | Y |
| | **Pil-Chp network** | | |
| *pilJ* | T570I | Methyl-accepting chemotaxis protein – transduces signal for cAMP synthesis | Y |
| *fimL* | L115F | Scaffold protein – involved in CyaB activation | Y |
| *chpA* | Δ36bp – 3776-3811 | Histidine kinase – required for cAMP upregulation | N |
| *chpA* | Δ39bp – 3856-3894 | - | N |
| | **Other** | | |
| *pilO* | P168L | Alignment complex structural protein | Y |
| *cpdA* | V258G | cAMP phosphodiesterase | Y |
| *cpdA* | 575-599 sequence duplication AGGTACGCTGCCTGCTGTGGGGGCA | - | N |
| *vfr* | G146R | Transcription factor | N |

**Table S2: All strains and plasmids used in this study.**

| Strains | | |
|---|---|---|
| **Strain name** | **Genotype** | **Source** |
| ***E. coli* strains** | | |
| DH5α | *F-φ80lacZΔM15 Δ(lacZYA-argF)U169 recA1 endA1 hsdR17(rk−, mk+) phoA supE44 thi-1 gyrA96 relA1 λ−* | Invitrogen |
| BL21-CodonPlus-RIL | *F– ompT hsdS (rB – mB – ) dcm+ Tetr gal endA Hte [argU ileY leuW Camr ]* | New England Biolabs |
| BTH101 | Bacterial 2-hybrid strain | Euromedex |
| SM10 | *thi-1 thr leu tonA lacY supE recA::RP4-2-Tc::Mu (KmR)* | Invitrogen |
| PilS/PilS | BTH101 with PilS in pUT18C and pKT25 vectors | (1) |
| T18/T25 | BTH101 with empty pUT18C and pKT25 vectors | This study |
| FimX/FimX | BTH101 with FimX in pUT18C and pKT25 vectors | This study |
| FimX AAA/FimX AAA | BTH101 with FimX AAA in pUT18C and pKT25 vectors | This study |
| FimX-T18/FimX AAA-T25 | BTH101 with FimX in pUT18C and FimX AAA in pKT25 vectors | This study |
| FimX-T25/FimX AAA-T18 | BTH101 with FimX AAA in pUT18C and FimX in pKT25 vectors | This study |
| FimX-T18/PilZ-T25 | BTH101 with FimX in pUT18C and PilZ in pKT25 vectors | This study |
| FimX-T25/PilZ-T18 | BTH101 with PilZ in pUT18C and FimX in pKT25 vectors | This study |
| FimX AAA-T18/PilZ-T25 | BTH101 with FimX AAA in pUT18C and PilZ in pKT25 vectors | This study |
| FimX AAA-T25/PilZ-T18 | BTH101 with PilZ in pUT18C and FimX AAA in pKT25 vectors | This study |
| ***P. aeruginosa* strains** | | |
| mPAO1 | WT | (2) |
| *pilA*::Tn5 | mPAO1 *pilA* transposon mutant | (2) |
| Δ*fimX* | mPAO1 *fimX* deletion | This study |
| FimX AAA | mPAO1 FimX EVL motif to AAA mutation | This study |
| Δ*pilZ* | mPAO1 *pilZ* deletion | This study |
| Δ*fimX pilZ* | mPAO1 *fimX pilZ* double deletion | This study |
| Δ*fimX pilB* Δ1bp | mPAO1 *fimX* deletion with *pilB* Δ1bp suppressor | This study |
| Δ*fimX* PilB T430P | mPAO1 *fimX* deletion with PilB T430P suppressor | This study |
| Δ*fimX* PilO P168L | mPAO1 *fimX* deletion with PilO P168L suppressor | This study |
| Δ*fimX* PilJ T750I | mPAO1 *fimX* deletion with PilJ T570I suppressor | This study |
| Δ*fimX* FimL L115F | mPAO1 *fimX* deletion with FimL L115F suppressor | This study |
| Δ*fimX* CpdA V258G | mPAO1 *fimX* deletion with CpdA V258G suppressor | This study |

| | | |
|---|---|---|
| mPAO1 + pMS402-*pcdrA*:LuX | mPAO1 with cdGMP reporter *pcdrA*:LuX cassette in pMS402 | This study |
| *pilA*::Tn5 + pMS402-*pcdrA*:LuX | *pilA*::Tn5 with cdGMP reporter *pcdrA*:LuX cassette | This study |
| mPAO1 + pMS402 | mPAO1 with empty pMS402 | This study |
| mPAO1 + pMS402-*pcdrA*:LuX pBADGr-*sadC* | mPAO1 with cdGMP reporter *pcdrA*:LuX cassette in pMS402 and *sadC* in pBADGr | This study |
| Δ*fimX* + pMS402-*pcdrA*:LuX | Δ*fimX* with cdGMP reporter *pcdrA*:LuX cassette in pMS402 | This study |
| Δ*fimX pilB* Δ1bp + pMS402-*pcdrA*:LuX | Δ*fimX pilB* Δ1bp with cdGMP reporter *pcdrA*:LuX cassette in pMS402 | This study |
| Δ*fimX* PilB T430P + pMS402-*pcdrA*:LuX | Δ*fimX* PilB T430P with cdGMP reporter *pcdrA*:LuX cassette in pMS402 | This study |
| Δ*fimX* PilO P168L + pMS402-*pcdrA*:LuX | Δ*fimX* PilO P168L with cdGMP reporter *pcdrA*:LuX cassette in pMS402 | This study |
| Δ*fimX* PilJ T750I + pMS402-*pcdrA*:LuX | Δ*fimX* PilJ T570I with cdGMP reporter *pcdrA*:LuX cassette in pMS402 | This study |
| Δ*fimX* FimL L115F + pMS402-*pcdrA*:LuX | Δ*fimX* L115F with cdGMP reporter *pcdrA*:LuX cassette in pMS402 | This study |
| Δ*fimX* CpdA V258G + pMS402-*pcdrA*:LuX | Δ*fimX* CpdA V258Gwith cdGMP reporter *pcdrA*:LuX cassette in pMS402 | This study |
| mPAO1 + pBADGr | mPAO1 with empty pBADGr | This study |
| mPAO1 + pBADGr-*sadC* | mPAO1 with *sadC* in pBADGr | This study |
| mPAO1 + pBADGr-*ydeH* | mPAO1 with *ydeH* in pBADGr | This study |
| Δ*fimX* + pBADGr | Δ*fimX* with empty pBADGr | This study |
| Δ*fimX* + pBADGr-*sadC* | Δ*fimX* with *sadC* in pBADGr | This study |
| Δ*fimX* + pBADGr-*ydeH* | Δ*fimX* with *ydeH* in pBADGr | This study |
| mPAO1 + pBADGr CyaB R456L | mPAO1 with CyaB R456L in pBADGr | (3) |
| mPAO1 + cAMP reporter | mPAO1 with cAMP reporter plasmid | This study |
| *pilA*::Tn5 + cAMP reporter | *pilA*::Tn5 with cAMP reporter plasmid | This study |
| Δ*cyaB* + cAMP reporter | Δ*cyaB* with cAMP reporter plasmid | This study |
| mPAO1 + cAMP reporter pBADGr-CyaB R456L | mPAO1 with cAMP reporter plasmid and CyaB R456L in pBADGr | This study |

| | | |
|---|---|---|
| Δ*fimX* + cAMP reporter | Δ*fimX* with cAMP reporter plasmid | This study |
| Δ*fimX pilB* Δ1bp + cAMP reporter | Δ*fimX pilB* Δ1bp with cAMP reporter plasmid | This study |
| Δ*fimX* PilB T430P + cAMP reporter | Δ*fimX* PilB T430P with cAMP reporter plasmid | This study |
| Δ*fimX* PilO P168L + cAMP reporter | Δ*fimX* PilO P168L with cAMP reporter plasmid | This study |
| Δ*fimX* PilJ T750I + cAMP reporter | Δ*fimX* PilJ T570I with cAMP reporter plasmid | This study |
| Δ*fimX* FimL L115F + cAMP reporter | Δ*fimX* FimL L115F with cAMP reporter plasmid | This study |
| Δ*fimX* CpdA V258G + cAMP reporter | Δ*fimX* CpdA V258G with cAMP reporter plasmid | This study |
| mPAO1 + pHERD30T | mPAO1 with empty pHERD30T | This study |
| *pilA*::Tn5 + pHERD30T | *pilA*::Tn5 with empty pHERD30T | This study |
| Δ*fimX* + pHERD30T | Δ*fimX* with empty pHERD30T | This study |
| Δ*fimX pilB* Δ1bp + pHERD30T | Δ*fimX pilB* Δ1bp with empty pHERD30T | This study |
| Δ*fimX* PilB T430P + pHERD30T | Δ*fimX* PilB T430P with empty pHERD30T | This study |
| Δ*fimX* PilO P168L + pHERD30T | Δ*fimX* PilO P168L with empty pHERD30T | This study |
| Δ*fimX* PilJ T750I + pHERD30T | Δ*fimX* PilJ T570I with empty pHERD30T | This study |
| Δ*fimX* FimL L115F + pHERD30T | Δ*fimX* FimL L115F with empty pHERD30T | This study |
| Δ*fimX* CpdA V258G + pHERD30T | Δ*fimX* CpdA V258G with empty pHERD30T | This study |
| mPAO1 + pHERD30T-*fimX* | mPAO1 with *fimX* in pHERD30T | This study |
| *pilA*::Tn5 + pHERD30T-*fimX* | *pilA*::Tn5 with *fimX* in pHERD30T | This study |
| Δ*fimX* + pHERD30T-*fimX* | Δ*fimX* with *fimX* in pHERD30T | This study |
| Δ*fimX pilB* Δ1bp + pHERD30T-*fimX* | Δ*fimX pilB* Δ1bp with *fimX* in pHERD30T | This study |
| Δ*fimX* PilB T430P + pHERD30T-*fimX* | Δ*fimX* PilB T430P with *fimX* in pHERD30T | This study |
| Δ*fimX* PilO P168L + pHERD30T-*fimX* | Δ*fimX* PilO P168L with *fimX* in pHERD30T | This study |
| Δ*fimX* PilJ T750I + pHERD30T-*fimX* | Δ*fimX* PilJ T570I with *fimX* in pHERD30T | This study |
| Δ*fimX* FimL L115F + pHERD30T-*fimX* | Δ*fimX* FimL L115F with *fimX* in pHERD30T | This study |
| Δ*fimX* CpdA V258G + pHERD30T-*fimX* | Δ*fimX* CpdA V258G with *fimX* in pHERD30T | This study |

| | | |
|---|---|---|
| mPAO1 + pHERD30T-FimX AAA | mPAO1 with FimX AAA pHERD30T | This study |
| *pilA*::Tn5 + pHERD30T-FimX AAA | *pilA*::Tn5 with FimX AAA pHERD30T | This study |
| Δ*fimX* + pHERD30T-FimX AAA | Δ*fimX* with FimX AAA pHERD30T | This study |
| Δ*fimX pilB* Δ1bp + pHERD30T-FimX AAA | Δ*fimX pilB* Δ1bp with FimX AAA pHERD30T | This study |
| Δ*fimX* PilB T430P + pHERD30T-FimX AAA | Δ*fimX* PilB T430P with FimX AAA pHERD30T | This study |
| Δ*fimX* PilO P168L + pHERD30T-FimX AAA | Δ*fimX* PilO P168L with FimX AAA pHERD30T | This study |
| Δ*fimX* PilJ T750I + pHERD30T-FimX AAA | Δ*fimX* PilJ T570I with FimX AAA pHERD30T | This study |
| Δ*fimX* FimL L115F + pHERD30T-FimX AAA | Δ*fimX* FimL L115F with FimX AAA pHERD30T | This study |
| Δ*fimX* CpdA V258G + pHERD30T-FimX AAA | Δ*fimX* CpdA V258G with FimX AAA pHERD30T | This study |
| Δ*fimX* + pHERD30T-$His_6$-CyaB | Δ*fimX* with $His_6$-CyaB in pHERD30T | This study |
| Δ*fimX* + pHERD30T-$His_6$-CyaB D234S | Δ*fimX* with $His_6$-CyaB D234S in pHERD30T | This study |
| Δ*fimX* Δ*cyaB* + pHERD30T | Δ*fimX* Δ*cyaB* with empty pHERD30T | This study |
| Δ*cyaB* + pHERD30T | Δ*cyaB* with empty pHERD30T | This study |
| Δ*cyaB* + pHERD30T-$His_6$-CyaB | Δ*cyaB* with $His_6$-CyaB in pHERD30T | This study |
| Δ*cyaB* + pHERD30T-$His_6$-CyaB D234S | Δ*cyaB* with $His_6$-CyaB D234S in pHERD30T | This study |
| Δ*pilG pilH* + pHERD30T | Δ*pilG pilH* with empty pHERD30T | This study |
| Δ*pilG pilH* + pHERD30T-$His_6$-CyaB | Δ*pilG pilH* with $His_6$-CyaB in pHERD30T | This study |
| Δ*pilG pilH* + pHERD30T-$His_6$-CyaB D234S | Δ*pilG pilH* with $His_6$-CyaB D234S in pHERD30T | This study |
| Δ*vfr* | mPAO1 *vfr* deletion | (4) |

| | | |
|---|---|---|
| *pilO*::FRT | mPAO1 *pilO* FRT insertion | (5) |
| mPAO1 *pvfr*:mRUBY3 | mPAO1 with mRUBY3 cassette inserted upstream of the *pilMNOPQ pvfr* promoter | This study |
| *pilA*::Tn5 *pvfr*:mRUBY3 | *pilA*::Tn5 with mRUBY3 cassette inserted upstream of the *pilMNOPQ pvfr* promoter | This study |
| Δ*fimX* *pvfr*:mRUBY3 | Δ*fimX* with mRUBY3 cassette inserted upstream of the *pilMNOPQ pvfr* promoter | This study |
| Δ*vfr pvfr*:mRUBY3 | Δ*vfr* with mRUBY3 cassette inserted upstream of the *pilMNOPQ pvfr* promoter | This study |
| Δ*cyaB* *pvfr*:mRUBY3 | Δ*cyaB* with mRUBY3 cassette inserted upstream of the *pilMNOPQ pvfr* promoter | This study |
| mPAO1 *pvfr*:mRUBY3 + pBADGr R456L | mPAO1 with mRUBY3 cassette inserted upstream of the *pilMNOPQ pvfr* promoter and CyaB R456L in pBADGr | This study |
| Δ*fimX pilB* Δ1bp *pvfr*:mRUBY3 | Δ*fimX pilB* Δ1bp with mRUBY3 cassette inserted upstream of the *pilMNOPQ pvfr* promoter | This study |
| Δ*fimX* PilB T430P *pvfr*:mRUBY3 | Δ*fimX* PilB T430P with mRUBY3 cassette inserted upstream of the *pilMNOPQ pvfr* promoter | This study |
| Δ*fimX* PilO P168L *pvfr*:mRUBY3 | Δ*fimX* PilO P168L with mRUBY3 cassette inserted upstream of the *pilMNOPQ pvfr* promoter | This study |
| Δ*fimX* PilJ T750I *pvfr*:mRUBY3 | Δ*fimX* PilJ T570I with mRUBY3 cassette inserted upstream of the *pilMNOPQ pvfr* promoter | This study |
| Δ*fimX* FimL L115F *pvfr*:mRUBY3 | Δ*fimX* FimL L115F with mRUBY3 cassette inserted upstream of the *pilMNOPQ pvfr* promoter | This study |
| Δ*fimX* CpdA V258G *pvfr*:mRUBY3 | Δ*fimX* CpdA V258G with mRUBY3 cassette inserted upstream of the *pilMNOPQ pvfr* promoter | This study |
| Δ*xcpT* | mPAO1 *xcpT* deletion | Burrows lab |
| Δ*fimX pilB*::Tn5 + pHERD30T | Δ*fimX pilB*::Tn5 with empty pHERD30T | This study |
| Δ*fimX pilB*::Tn5 + pHERD30T-*pilB* | Δ*fimX pilB*::Tn5 with *pilB* in pHERD30T | This study |
| Δ*fimX pilB*::Tn5 + pHERD30T-*pilB* Δ1bp | Δ*fimX pilB*::Tn5 with *pilB* Δ1bp in pHERD30T | This study |
| Δ*fimX pilB*::Tn5 + pHERD30T-*pilB* Δ1bp-3 | Δ*fimX pilB*::Tn5 with *pilB* Δ1bp-3 in pHERD30T | This study |
| Δ*fimX pilB*::Tn5 + pHERD30T-*pilB* Δ1bp-6 | Δ*fimX pilB*::Tn5 with *pilB* Δ1bp-6 in pHERD30T | This study |
| Δ*fimX pilB*::Tn5 + pHERD30T-*pilB* Δ1bp-9 | Δ*fimX pilB*::Tn5 with *pilB* Δ1bp-9 in pHERD30T | This study |
| Δ*fimX pilB*::Tn5 + pHERD30T-*pilB* Δ1bp-12 | Δ*fimX pilB*::Tn5 with *pilB* Δ1bp-12 in pHERD30T | This study |
| Δ*fimX pilB*::Tn5 + pHERD30T-*pilB* Δ1bp-15 | Δ*fimX pilB*::Tn5 with *pilB* Δ1bp-15 in pHERD30T | This study |
| Δ*fimX pilB*::Tn5 + pHERD30T-*pilB* Δ1bp-18 | Δ*fimX pilB*::Tn5 with *pilB* Δ1bp-18 in pHERD30T | This study |

| | | |
|---|---|---|
| Δ*fimX pilB*::Tn5 + pHERD30T-*pilB* Δ1bp-21 | Δ*fimX pilB*::Tn5 with *pilB* Δ1bp-21 in pHERD30T | This study |
| *pilB*::Tn5 | mPAO1 *pilB* transposon mutant | (2) |
| *pilB*::Tn5 + pHERD30T | *pilB*::Tn5 with empty pHERD30T | This study |
| *pilB*::Tn5 + pHERD30T-*pilB* | *pilB*::Tn5 with *pilB* in pHERD30T | This study |
| *pilB*::Tn5 + pHERD30T-*pilB* Δ1bp | *pilB*::Tn5 with *pilB* Δ1bp in pHERD30T | This study |
| *pilB*::Tn5 + pHERD30T-*pilB* Δ1bp-3 | *pilB*::Tn5 with *pilB* Δ1bp-3 in pHERD30T | This study |
| *pilB*::Tn5 + pHERD30T-*pilB* Δ1bp-6 | *pilB*::Tn5 with *pilB* Δ1bp-6 in pHERD30T | This study |
| *pilB*::Tn5 + pHERD30T-*pilB* Δ1bp-9 | *pilB*::Tn5 with *pilB* Δ1bp-9 in pHERD30T | This study |
| *pilB*::Tn5 + pHERD30T-*pilB* Δ1bp-12 | *pilB*::Tn5 with *pilB* Δ1bp-12 in pHERD30T | This study |
| *pilB*::Tn5 + pHERD30T-*pilB* Δ1bp-15 | *pilB*::Tn5 with *pilB* Δ1bp-15 in pHERD30T | This study |
| *pilB*::Tn5 + pHERD30T-*pilB* Δ1bp-18 | *pilB*::Tn5 with *pilB* Δ1bp-18 in pHERD30T | This study |
| *pilB*::Tn5 + pHERD30T-*pilB* Δ1bp-21 | *pilB*::Tn5 with *pilB* Δ1bp-21 in pHERD30T | This study |
| mPAO1 + pHERD30T-*pilB* | mPAO1 with *pilB* in pHERD30T | This study |
| mPAO1 + pHERD30T-*pilB* Δ1bp | mPAO1 with *pilB* Δ1bp in pHERD30T | This study |
| mPAO1 + pHERD30T-*pilB* Δ1bp-3 | mPAO1 with *pilB* Δ1bp-3 in pHERD30T | This study |
| mPAO1 + pHERD30T-*pilB* Δ1bp-6 | mPAO1 with *pilB* Δ1bp-6 in pHERD30T | This study |
| mPAO1 + pHERD30T-*pilB* Δ1bp-9 | mPAO1 with *pilB* Δ1bp-9 in pHERD30T | This study |
| mPAO1 + pHERD30T-*pilB* Δ1bp-12 | mPAO1 with *pilB* Δ1bp-12 in pHERD30T | This study |
| mPAO1 + pHERD30T-*pilB* Δ1bp-15 | mPAO1 with *pilB* Δ1bp-15 in pHERD30T | This study |

| | | |
|---|---|---|
| mPAO1 + pHERD30T-*pilB* Δ1bp-18 | mPAO1 with *pilB* Δ1bp-18 in pHERD30T | This study |
| mPAO1 + pHERD30T-*pilB* Δ1bp-21 | mPAO1 with *pilB* Δ1bp-21 in pHERD30T | This study |
| mPAO1 PilB D566V | PilB D566V chromosomal knock-in | This study |
| mPAO1 PilB D566K | PilB D566K chromosomal knock-in | This study |
| mPAO1 PilB D566E | PilB D566E chromosomal knock-in | This study |
| mPAO1 PilB $D566^{STOP}$ | PilB D566 to STOP codon chromosomal knock-in | This study |
| Δ*fimX pilB*::Tn5 + PilB T430P | Δ*fimX pilB*::Tn5 with PilB T430P in pHERD30T | This study |
| Δ*fimX pilB*::Tn5 + PilB T430S | Δ*fimX pilB*::Tn5 with PilB T430S in pHERD30T | This study |
| Δ*fimX pilB*::Tn5 + PilB T430N | Δ*fimX pilB*::Tn5 with PilB T430N in pHERD30T | This study |
| Δ*fimX pilB*::Tn5 + PilB T430I | Δ*fimX pilB*::Tn5 with PilB T430I in pHERD30T | This study |
| Δ*fimX pilB*::Tn5 + PilB L429P | Δ*fimX pilB*::Tn5 with PilB L429P in pHERD30T | This study |
| Δ*fimX pilB*::Tn5 + PilB R431P | Δ*fimX pilB*::Tn5 with PilB R431P in pHERD30T | This study |
| Δ*fimX pilB*::Tn5 + PilB R431A | Δ*fimX pilB*::Tn5 with PilB R431A in pHERD30T | This study |
| *pilB*::Tn5 + PilB T430P | *pilB*::Tn5 with PilB T430P in pHERD30T | This study |
| *pilB*::Tn5 + PilB T430S | *pilB*::Tn5 with PilB T430S in pHERD30T | This study |
| *pilB*::Tn5 + PilB T430N | *pilB*::Tn5 with PilB T430N in pHERD30T | This study |
| *pilB*::Tn5 + PilB T430I | *pilB*::Tn5 with PilB T430I in pHERD30T | This study |
| *pilB*::Tn5 + PilB L429P | *pilB*::Tn5 with PilB L429P in pHERD30T | This study |
| *pilB*::Tn5 + PilB R431P | *pilB*::Tn5 with PilB R431P in pHERD30T | This study |
| *pilB*::Tn5 + PilB R431A | Δ*pilB*::Tn5 with PilB R431A in pHERD30T | This study |
| mPAO1 + PilB T430P | mPAO1 with PilB T430P in pHERD30T | This study |
| mPAO1 + PilB T430S | mPAO1with PilB T430S in pHERD30T | This study |
| mPAO1 + PilB T430N | mPAO1Tn5 with PilB T430N in pHERD30T | This study |
| mPAO1 + PilB T430I | mPAO1 with PilB T430I in pHERD30T | This study |
| mPAO1 + PilB L429P | mPAO1 with PilB L429P in pHERD30T | This study |
| mPAO1 + PilB R431P | mPAO1 with PilB R431P in pHERD30T | This study |

| | | |
|---|---|---|
| *pilB*::Tn5 + PilB R431A | mPAO1 with PilB R431A in pHERD30T | This study |
| *pilB* Δ1bp | mPAO1 *pilB* Δ1bp chromosomal knock-in | This study |
| PilB T430P | mPAO1 PilB T430P chromosomal knock-in | This study |
| *pilB* Δ1bp FimX AAA | *pilB* Δ1bp FimX AAA chromosomal knock-in | This study |
| PilB T430P FimX AAA | PilB T430P FimX AAA chromosomal knock-in | This study |
| PilA A86C | PilA A86C chromosomal knock-in | This study |
| Δ*fimX* PilA A86C | Δ*fimX* PilA A86C chromosomal knock-in | This study |
| Δ*fimX pilB* Δ1bp PilA A86C | Δ*fimX pilB* Δ1bp PilA A86C chromosomal knock-in | This study |
| *pilB* Δ1bp PilA A86C | *pilB* Δ1bp PilA A86C chromosomal knock-in | This study |
| Δ*fimX* CpdA V258G PilA A86C | Δ*fimX* CpdA V258G PilA A86C chromosomal knock-in | This study |
| Δ*pilT* | mPAO1 *pilT* deletion | This study |
| PilA A86C Δ*pilT* | PilA A86C chromosomal knock-in and *pilT* deletion | This study |
| Δ*fimX* PilA A86C Δ*pilT* | Δ*fimX* PilA A86C chromosomal knock-in and *pilT* deletion | This study |
| Δ*fimX pilB* Δ1bp PilA A86C Δ*pilT* | Δ*fimX pilB* Δ1bp PilA A86C chromosomal knock-in and *pilT* deletion | This study |
| *pilB* Δ1bp PilA A86C Δ*pilT* | *pilB* Δ1bp PilA A86C chromosomal knock-in and *pilT* deletion | This study |
| Δ*fimX* CpdA V258G PilA A86C Δ*pilT* | Δ*fimX* CpdA V258G PilA A86C chromosomal knock-in and *pilT* deletion | This study |
| Δ*fimX* + pHERD30T-mNGr-FimX | Δ*fimX* with FimX N-terminally tagged with mNeonGreen in pHERD30T | This study |
| Δ*fimX pilB*::Tn5 + pHERD30T-mNGr-FimX | Δ*fimX pilB*::Tn5 with FimX N-terminally tagged with mNeonGreen in pHERD30T | This study |
| Δ*fimX* + pHERD30T-mNGr-FimX AAA | Δ*fimX* with FimX AAA N-terminally tagged with mNeonGreen in pHERD30T | This study |
| Δ*fimX pilB*::Tn5 + pHERD30T-mNGr-FimX AAA | Δ*fimX pilB*::Tn5 with FimX AAA N-terminally tagged with mNeonGreen in pHERD30T | This study |
| **Plasmid list** | | |
| **Plasmid name** | **Characteristics** | **Source** |
| pEX18Gm | Suicide vector for gene replacement | (6) |
| pEX18Gm-Δ*fimX* | *fimX* deletion construct | This study |
| pEX18Gm-Δ*pilZ* | *pilZ* deletion construct | This study |
| pEX18Gm-Δ*cyaB* | *cyaB* deletion construct | This study |
| pEX18Gm-*pilB* Δ1bp | *pilB* Δ1bp (A1697) deletion construct | This study |

| | | |
|---|---|---|
| pEX18Gm-PilB T430P | PilB T430P chromosomal knock-in construct | This study |
| pEX18Gm-CpdA V258G | CpdA V258G chromosomal knock-in construct | This study |
| pEX18Gm- Δ*pilG pilH* | *pilG* and *pilH* double deletion construct | This study |
| pEX18Gm-*pvfr*-mRUBY3 | mRUBY3 knock-in construct upstream of *pilMNOPQ vfr* promoter | This study |
| pEX18Gm-PilB D566V | PilB D566V chromosomal knock-in construct | This study |
| pEX18Gm-PilB D566K | PilB D566K chromosomal knock-in construct | This study |
| pEX18Gm-PilB D566E | PilB D566E chromosomal knock-in construct | This study |
| pEX18Gm-PilB $D566^{STOP}$ | PilB D566 STOP codon chromosomal knock-in construct | This study |
| pEX18Gm-FimX AAA | FimX EVL motif to AAA chromosomal knock-in construct | This study |
| pEX18Gm-PilA A86C | PilA A86C chromosomal knock-in construct | This study |
| pEX18Gm-Δ*pilT* | *pilT* deletion construct | Burrows lab |
| pHERD30T | Broad host-range expression vector | (7) |
| pHERD30T-*fimX* | *fimX* expression construct | This study |
| pHERD30T-FimX AAA | FimX AAA expression construct | This study |
| pHERD30T-$His_6$-CyaB | $His_6$-CyaB expression construct | This study |
| pHERD30T-$His_6$-CyaB D234S | $His_6$-CyaB D234S expression construct | This study |
| pHERD30T-*pilB* | PilB expression construct | This study |
| pHERD30T-*pilB* Δ1bp | *pilB* Δ1bp expression construct | This study |
| pHERD30T-*pilB* Δ1bp-3 | *pilB* Δ1bp with one codon deleted from the 3′ end expression construct | This study |
| pHERD30T-*pilB* Δ1bp-6 | *pilB* Δ1bp with two codons deleted from the 3′ end expression construct | This study |
| pHERD30T-*pilB* Δ1bp-9 | *pilB* Δ1bp with three codons deleted from the 3′ end expression construct | This study |
| pHERD30T-*pilB* Δ1bp-12 | *pilB* Δ1bp with four codons deleted from the 3′ end expression construct | This study |
| pHERD30T-*pilB* Δ1bp-15 | *pilB* Δ1bp with five codons deleted from the 3′ end expression construct | This study |
| pHERD30T-*pilB* Δ1bp-18 | *pilB* Δ1bp with six codons deleted from the 3′ end expression construct | This study |
| pHERD30T-*pilB* Δ1bp-21 | *pilB* Δ1bp with seven codons deleted from the 3′ end expression construct | This study |
| pHERD30T-PilB T430P | PilB T430P expression construct | This study |
| pHERD30T-PilB T430S | PilB T430S expression construct | This study |
| pHERD30T-PilB T430N | PilB T430N expression construct | This study |

| | | |
|---|---|---|
| pHERD30T-PilB T430I | PilB T430I expression construct | This study |
| pHERD30T-PilB L429P | PilB L429P expression construct | This study |
| pHERD30T-PilB R431P | PilB R431P expression construct | This study |
| pHERD30T-PilB R431A | PilB R431A expression construct | This study |
| pHERD30T-mNGr-FimX | FimX with N-terminal mNeonGreen fusion expression construct | This study |
| pHERD30T-mNGr-FimX AAA | FimX AAA with N-terminal mNeonGreen fusion expression construct | This study |
| pBADGr | Broad host-range expression vector | (8) |
| pBADGr-*sadC* | *sadC* expression construct | (4) |
| pBADGr-*ydeH* | *ydeH* expression construct | Howell lab |
| pBADGr-CyaB R456L | CyaB R456L expression construct | (3) |
| pMS402 | Broad host-range expression vector | (4) |
| pMS402-*pcdrA*:LuX | cdGMP reporter construct | (4) |
| cAMP reporter (*PaQa*:YFP/*rpoD*: mRUBY3) | cAMP reporter plasmid | (9) |
| pET28b-ΔN1D-$His_6$-PilB | N-terminally truncated $His_6$-tagged PilB for heterologous expression in *E. coli* | This study |
| pET28b-ΔN1D-$His_6$-PilB K332A | N-terminally truncated $His_6$-tagged PilB K332A for heterologous expression in *E. coli* | This study |
| pET28b-ΔN1D-$His_6$-*PilB* Δ1bp | N-terminally truncated $His_6$-tagged *pilB* Δ1bp for heterologous expression in *E. coli* | This study |
| pET28b-ΔN1D-$His_6$-PilB T430P | N-terminally truncated $His_6$-tagged PilB T430P for heterologous expression in *E. coli* | This study |
| pET28b-$His_6$-FimX | $His_6$-tagged FimX for heterologous expression in *E. coli* | This study |

### Table S3: Primers used in this study.

| Primers | | |
|---|---|---|
| **Primer name** | **Characteristics** | **Sequence** |
| ΔFimX UpS_Fwd | For making delta FimX construct - Upstream region | ATATGAGCTCACGCAGATGGGCATCGAG |
| ΔFimX UpS_Rev | For making delta FimX construct - Upstream region | ATATTCTAGAATGTACGCGGGTCGCGT |
| ΔFimX DnS_Fwd | For making delta FimX construct - Downstream region | ATATTCTAGAGAGAGCGCCAGCGTCCTC |
| ΔFimX DnS_Rev | For making delta FimX construct - Downstream region | ATATAAGCTTCAACGAGTTGCAGATCGTCG AC |
| ΔPilZ UpS_Fwd | For making delta PilZ construct - Upstream region | ATATGAATTCGAGGCCCTGGAGGAACTGTT |
| ΔPilZ UpS_Rev | For making delta PilZ construct - Upstream region | ATATTCTAGACAGATTGGGTGGCAAACTCA T |
| ΔPilZ DnS_Fwd | For making delta PilZ construct - Downstream region | ATATTCTAGAAGTTCAACGACGGTGACAAC AC |
| ΔPilZ DnS_Rev | For making delta PilZ construct - Downstream region | ATATAAGCTTGTGCACGATCACTGGCTTG |
| cpdA UpS_fwd | For making delta CpdA construct - Upstream region | ATATGAATTCAAGGGGAGCGGCTGCTC |
| cpdA UpS_rev | For making delta CpdA construct - Upstream region | ATATTCTAGATGGCTGTCCGAGAGCTGC |
| cpdA DnS_fwd | For making delta CpdA construct - Downstream region | ATATTCTAGACCTGGAAACGGGGATCTCG |
| cpdA DnS_rev | For making delta CpdA construct - Downstream region | ATATAAGCTTGTGGTCCTCGGTGAGTTCCC |
| cpdA V258G_fwd | For making V258G knockin construct when amplified from ΔfimX HT genomic template - pairs with cpdA DnS_rev primer | ATATGAATTCGGTTCGCCGGTAACCACG |
| FimX pHERD_Fwd | For cloning PAO1 FimX into pHERD | ATATTCTAGACTGAGCCCTTTCCATGG |
| FimX pHERD_Rev | For cloning PAO1 FimX into pHERD | ATATAAGCTTTCATTCGTCTCCCGAGG |
| PilZ pHERD_Fwd | For cloning PAO1 PilZ into pHERD | ATATGAATTCGGCAGGAACCTGCATGA |
| PilZ pHERD_Rev | For cloning PAO1 PilZ into pHERD | ATATAAGCTTTTACATCGTGTGGGTCGG |
| CpdA pHERD_Fwd | For cloning PAO1 cpdA into pHERD | ATATGAATTCAGGAGACGGCCCCTTG |
| CpdA pHERD_Rev | For cloning PAO1 cpdA into pHERD | ATATAAGCTTTCAGTATCCGGCGGTGT |
| CyaB UpS | For cloning CyaB upstream region | ATATGAATTCGAGTTCTACCCCTACTACCTG CAG |
| CyaB UpS_Rev | For cloning CyaB upstream region | ATATTCTAGACACGCGCGAATAGTATTCAC |
| CyaB DnS_Fwd | For cloning CyaB downstream region | ATATTCTAGAACTACGACAAGGAACGGGTC |
| CyaB DnS_Rev | For cloning CyaB downstream region | ATATAAGCTTAAAAGAACCTGGAGGCGTTC |
| FimX AAA_Fwd | For introducing AAA at the FimX EVL motif - has NotI cut site | GCCACGAGAACTACGCGGCCGCCCTGCGCC TGCTCAAT |

| | | |
|---|---|---|
| FimX AAA_Rev | For introducing AAA at the FimX EVL motif - has NotI cut site | ATTGAGCAGGCGCAGGGCGGCCGCGTAGTTCTCGTGGC |
| PilB pHERD_Fwd | For cloning PilB into pHERD30T | ATATGAGCTCGCGATTCCTTCCCCATGA |
| PilB WT pHERD_Rev | For cloning wild-type PilB into pHERD30T | ATATTCTAGATTAATCCTTGGTCACGCGG |
| PilB d1bp pHERD | For cloning pilB delta 1bp into pHERD30T: pairs with PilB pHERD_Fwd | ATATTCTAGATTAACGCTTTGTCCGCCAT |
| PilB d1bp pHERD-3 | For cloning pilB delta 1bp into pHERD30T: pairs with PilB pHERD_Fwd | ATATTCTAGATTACTTTGTCCGCCATGGATTAAC |
| PilB d1bp pHERD-6 | For cloning pilB delta 1bp into pHERD30T: pairs with PilB pHERD_Fwd | ATATTCTAGATTATGTCCGCCATGGATTAACCT |
| PilB d1bp pHERD-9 New | For cloning pilB delta 1bp into pHERD30T: pairs with PilB pHERD_Fwd | ATATTCTAGATTACCGCCATGGATTAACCTTG |
| PilB d1bp pHERD-12 New | For cloning pilB delta 1bp into pHERD30T: pairs with PilB pHERD_Fwd | ATATTCTAGATTACCATGGATTAACCTTGGTCACG |
| PilB d1bp pHERD-15 New | For cloning pilB delta 1bp into pHERD30T: pairs with PilB pHERD_Fwd | ATATTCTAGATTATGGATTAACCTTGGTCACGC |
| PilB d1bp pHERD-18 New | For cloning pilB delta 1bp into pHERD30T: pairs with PilB pHERD_Fwd | ATATTCTAGATTAATTAACCTTGGTCACGCGGTT |
| PilB d1bp pHERD-21 New | For cloning pilB delta 1bp into pHERD30T: pairs with PilB pHERD_Fwd | ATATTCTAGATTAAACCTTGGTCACGCGGTT |
| PilM UpS-vfr rp_F | For cloning PilM upstream region into pEX18Gm to make vfr promoter mRUBY3 reporter | ATATGAATTCGCATTAGGCTTTTCACATCGAC |
| PilM UpS-vfr rp_R2 | For cloning PilM upstream region into pEX18Gm to make vfr promoter mRUBY3 reporter | ATATGGTACCTTCCCTATTAGCGTTCAATACTTACG |
| mRUBY3-vfr rp_F New2 | For cloning mRUBY3 into pEX18Gm to make vfr promoter reporter | ATATGGTACCATGGTGTCTAAGGGCGAAGAGC |
| mRUBY3-vfr rp_R New | For cloning mRUBY3 into pEX18Gm to make vfr promoter reporter | ATATTCTAGATTACTTGTACAGCTCGTCCATGCC |
| PilM DnS-vfr rp_F | For cloning PilM downstream region into pEX18Gm to make vfr promoter mRUBY3 reporter | ATATTCTAGAGTGCTAGGGCTCATAAAGAAGAAAG |
| PilM DnS-vfr rp_R | For cloning PilM downstream region into pEX18Gm to make vfr promoter mRUBY3 reporter | ATATAAGCTTCTGCTCAGCAGCGCATAG |
| CyaB D234S_F | For mutating CyaB D234 to S (has XhoI cut site): Pairs with CyaB pHERD_Rev | ACCGTGTTCTTCTCGAGCATCCGCGGCTTCACCGAG |
| CyaB D234S_R | For mutating CyaB D234 to S (has XhoI cut site): Pairs with CyaB pHERD_Fwd | CTCGGTGAAGCCGCGGATGCTCGAGAAGAACACGGT |

| | | |
|---|---|---|
| CyaB His_Fwd | For cloning CyaB with N-terminal His-tag into pHERD30T - encodes his tag | ATATGAATTCATGCACCACCACCACCACCA CATGAAGCCTACCCTCCCCG |
| CyaB His_Rev | For cloning CyaB with N-terminal His-tag into pHERD30T | ATATAAGCTTTTAGAGGATGACCTTGTCGC G |
| PilB N2D/CTD_Fwd | Encodes Pa PilB N2D and CTD with ΔD180-N terminus truncation for cloning into pET28b : Pairs with: Pa PilB_Rev | ATATGCTAGCGACGCACCTGTAGTACGTTT CGTC |
| Pa PilB_Rev | For cloning PilB into pET28b | ATATGAATTCTTAATCCTTGGTCACGCGGTT |
| Pa PilB d1bp pET28_Rev | For cloning PilB d1bp into pET28b. Pairs with PilB N2D/CTD_Fwd | ATATGAATTCTTAACGCTTTGTCCGCCATG |
| Pa PilB K332A_Fwd | For mutating PilB Walker A motif K332 to A | CCCACCGGCTCGGGCGCGACGGTATCGCTA TACACC |
| Pa PilB K332A_Rev | For mutating PilB Walker A motif K332 to A | GGTGTATAGCGATACCGTCGCGCCCGAGCC GGTGGG |
| PilB T430P_Fwd | For knocking in PilB T430 to P forward mutagenesis primer | GCCGCCGAGACCCTGCCCCGGTTGCTGAAC ATGG |
| PilB T430P_Rev | For knocking in PilB T430 to P reverse mutagenesis primer | CCATGTTCAGCAACCGGGGCAGGGTCTCGG CGGC |
| PilB T430N_F | For mutating PilB T430 to N forward primer | AGCGCCGCCGAGACCCTGAACCGGTTGCTG AACATGGGC |
| PilB T430N_R | For mutating PilB T430 to N reverse primer | GCCCATGTTCAGCAACCGGTTCAGGGTCTC GGCGGCGCT |
| PilB T430S_F | For mutating PilB T430 to S forward primer | AGCGCCGCCGAGACCCTGTCCCGGTTGCTG AACATGGGC |
| PilB T430S_R | For mutating PilB T430 to S reverse primer | GCCCATGTTCAGCAACCGGGACAGGGTCTC GGCGGCGCT |
| PilB T430I_F | For mutating PilB T430 to I forward primer | AGCGCCGCCGAGACCCTGATCCGGTTGCTG AACATGGGC |
| PilB T430I_R | For mutating PilB T430 to I reverse primer | GCCCATGTTCAGCAACCGGATCAGGGTCTC GGCGGCGCT |
| PilB R431A_F | For mutating PilB R431 to A forward primer | AGCGCCGCCGAGACCCTGACCGCGTTGCTG AACATGGGC |
| PilB R431A_R | For mutating PilB R431 to A reverse primer | GCCCATGTTCAGCAACGCGGTCAGGGTCTC GGCGGCGCT |
| PilB D566X UpS_Fwd | For knocking in PilB C-terminal residue upstream region: pairs with PilB D566X DnS_Rev | ATATGAATTCAGATCCGCGACCTGGAGAC |
| PilB D566X DnS_Rev | For knocking in PilB C-terminal residue upstream region: pairs with PilB D566X UpS_Fwd | ATATTCTAGACGATTCCGTTTTTCCTTGTA GGT |
| PilB D566V_Fwd | For knocking in PilB D566 to V forward mutagenesis primer - removes AseI cut site | AACCGCGTGACCAAGGTTTAATCCATGGCG GACA |
| PilB D566V_Rev | For knocking in PilB D566 to V reverse mutagenesis primer - removes AseI cut site | TGTCCGCCATGGATTAAACCTTGGTCACGC GGTT |
| PilB D566K_Fwd | For knocking in PilB D566 to K forward mutagenesis primer - removes AseI cut site | AACCGCGTGACCAAGAAGTAATCCATGGCG GACA |
| PilB D566K_Rev | For knocking in PilB D566 to K reverse mutagenesis primer - removes AseI cut site | TGTCCGCCATGGATTACTTCTTGGTCACGCG GTT |

| | | |
|---|---|---|
| PilB D566E_Fwd | For knocking in PilB D566 to E forward mutagenesis primer - removes AseI cut site | AACCGCGTGACCAAGGAGTAATCCATGGCGGACA |
| PilB D566E_Rev | For knocking in PilB D566 to E reverse mutagenesis primer - removes AseI cut site | TGTCCGCCATGGATTACTCCTTGGTCACGCGGTT |
| PilB D566STOP_Fwd | For knocking in PilB D566 to STOP codon forward mutagenesis primer - removes AseI cut site | AACCGCGTGACCAAGTAATAATCCATGGCGGACA |
| PilB D566STOP_Rev | For knocking in PilB D566 to STOP codon reverse mutagenesis primer - removes AseI cut site | TGTCCGCCATGGATTATTACTTGGTCACGCGGTT |
| PilB T430P UpS_Fwd | For knocking in PilB T430 to P forward mutagenesis primer - removes natural EcoRV cut site | ATATGAATTCGATATATCCGAACGACGCAAAC |
| PilB T430P DnS_Rev | For knocking in PilB T430 to P reverse mutagenesis primer | ATATTCTAGACTGCCCGGTCAGTCAGTTC |
| PilB T430P_Fwd | For knocking in PilB T430 to P forward mutagenesis primer | GCCGCCGAGACCCTGCCCCGGTTGCTGAACATGG |
| PilB T430P_Rev | For knocking in PilB T430 to P reverse mutagenesis primer | CCATGTTCAGCAACCGGGGCAGGGTCTCGGCGGC |
| FimX pET28_Fwd | For cloning FimX into pet28b | ATATGCTAGCATGGCCATCGAAAAGAAAACC |
| FimX pET28_Rev | For cloning FimX into pet28b | ATATAAGCTTTTATTCGTCTCCCGAGGAG |
| PilA A86C Fwd | For mutating PAO1 PilA A86C | GGCGTCGAGCCGGATTGTAACAAGTTGGGTGTA |
| PilA A86C Rev | For mutating PAO1 PilA A86C | TACACCCAACTTGTTACAATCCGGCTCGACGCC |
| PIlA A86 Ups_Fwd | PilA A86 primer for upstream region amplification | ATATGAATTCCGCTCAGTTGGATGCTGTC |
| PIlA A86 Dns_Rev | PilA A86 primer for downstream region amplification | ATATTCTAGAGCCAAGCTGGAAGCTTCC |
| PilG Ups_Fwd | For making a PilG and PilH double deletion | ATATGAGCTCTCCGCTTCCAGTTCGAAC |
| PilG Ups_Rev | For making a PilG and PilH double deletion | ATATTCTAGAGAATCGTTTTCGAATCGTCG |
| PilH Dns_Fwd | For making a PilG and PilH double deletion | ATATTCTAGATGGACGAAGAGACCCTGC |
| PilH Dns_Rev | For making a PilG and PilH double deletion | ATATAAGCTTAGCTGTTCGCTGAAGGTGTC |
| mNeonGreen-FimX_F1 | For making an mNeonGreen-FimX N-terminal fusion construct - has FimX RBS and adds ATG start codon | ATATTCTAGACTGAGCCCTTTCCATGGTGAGCAAGGGCGAGGAG |
| mNeonGreen-FimX_R1 | For making an mNeonGreen-FimX N-terminal fusion construct - encodes 5G linker | ACCACCACCACCACCCTTGTACAGCTCGTCCATGCC |
| mNeonGreen-FimX_F2 | For making an mNeonGreen-FimX N-terminal fusion construct - encodes 5G linker | GGTGGTGGTGGTGGTATGGCCATCGAAAAGAAAACC |
| mNeonGreen-FimX_R2 | For making an mNeonGreen-FimX N-terminal fusion construct | ATATAAGCTTTCATTCGTCTCCCGAGGAGA |
| PilB R431P_F | For mutating PilB R431 to P. Pairs with: PilB WT pHERD_Rev | AGCGCCGCCGAGACCCTGACCCCGTTGCTGAACATGGGC |

| | | |
|---|---|---|
| PilB R431P_R | For mutating PilB R431 to P. Pairs with:PilB pHERD_Fwd | GCCCATGTTCAGCAACGGGGTCAGGGTCTC GGCGGCGCT |
| PilB L429P_F | For mutating PilB L429 to P. Pairs with: PilB WT pHERD_Rev | AGCGCCGCCGAGACCCCGACCCGGTTGCTG AACATGGGC |
| PilB L429P_R | For mutating PilB L429 to P. Pairs with:PilB pHERD_Fwd | GCCCATGTTCAGCAACCGGGTCGGGGTCTC GGCGGCGCT |
